# Loss of SUMOylation drives aberrant PRC1 clustering and 3D genome rewiring independent of H3K27me3

**DOI:** 10.64898/2026.02.05.704038

**Authors:** Nazli Akilli, Paul-Swann Puel, Marco Di Stefano, Fernando Muzzopappa, Lauriane Fritsch, Fabien Erdel, Daniel Jost, Thierry Cheutin, Giacomo Cavalli

**Affiliations:** IGH, UMR9002, University of Montpellier, CNRS, INSERM, 141 Rue de la Cardonille, 34396, Montpellier Cedex 5, France; Laboratoire de Biologie et Modélisation de la Cellule, École Normale Supérieure de Lyon,CNRS, UMR5239, Inserm U1293, Université Claude Bernard Lyon 1; MCD, Center for Integrative Biology (CBI), University of Toulouse, CNRS, Toulouse, France

## Abstract

Polycomb Repressive Complex 1 (PRC1) forms nuclear condensates that organize target chromatin domains. SUMOylation modulates PRC1 clustering, but its impact on condensate properties and 3D genome architecture remains unclear. Here, we show that depletion of SUMO in *Drosophila* wing imaginal discs transforms PRC1 condensates into large structures with reduced molecular dynamics. Strikingly, this biophysical reorganization occurs without global loss of the H3K27me3 mark. Instead, Hi-C reveals widespread rewiring of topologically associating domain (TAD) interactions. PRC1-bound TADs lose specific long-range contacts with each other while gaining ectopic interactions with active chromatin. These topological shifts correlate with gene misregulation independently of changes in canonical Polycomb histone modifications. Our results establish SUMOylation as a critical regulator of PRC1 condensates, demonstrating that post-translational control of biomolecular condensation dictates 3D genome architecture and transcriptional output through mechanisms separable from histone mark deposition.

## Introduction

Polycomb group (PcG) proteins are evolutionarily conserved chromatin regulators that establish transcriptional silencing through the formation of repressive chromatin compartments. Canonical Polycomb repressive complex 1 (cPRC1), composed of core proteins Pc, Ph, Psc and Sce, contributes to chromatin compaction, histone H2A ubiquitylation, and long-range chromatin interactions that coordinate gene repression^1^. Recent studies have revealed that PRC1 can assemble into nuclear condensates, potentially via phase separation, thereby influencing genome topology and transcriptional memory^2–6^. PRC1 condensates coincide with clustered genomic targets, highlighting a close link between condensate formation and PRC1-mediated chromatin organization^7–9^.

In vitro studies on reconstituted nucleosome arrays have shown that both *Drosophila* and mammalian PRC1 can compact chromatin into dense aggregates that compete remodeling by Swi/Snf complexes^10^. Chromatin compaction is mediated by intrinsically disordered regions (IDRs) of PRC1 subunits, including Psc in *Drosophila* and CBX2 in mammalian cPRC1^1,11–13^. While these protein domains are central to phase separation, PRC1 is also subject to post-translational modifications such as glycosylation and SUMOylation, which can regulate the dynamics of membraneless nuclear condensates^14–16^. However, the contribution of such modifications to PRC1 phase behavior remains poorly understood.

SUMOylation involves the covalent attachment of a small ubiquitin-like modifier (SUMO) polypeptide to lysine residues of target proteins, altering their interactions and nuclear organization^17–18^. Pc, a core subunit of PRC1, has been shown to be SUMOylated in the nucleus, and reduced SUMOylation leads to the formation of unusually large Pc foci of ∼1 μm compared with less than 500 nm under normal conditions^15^. Similarly, the PRC1 subunit Scm is SUMOylated, and perturbation of SUMO levels induces Scm-dependent changes in target gene expression^14^. Despite these observations, the precise contribution of SUMOylation to PRC1 phase behavior and the resulting impact on chromatin architecture and transcriptional outcomes remains unclear.

Here, we investigate how integrity of the nuclear SUMOylation pathway modulates the phase behavior and functional output of PRC1. We hypothesized that if hypo-SUMOylated PRC1 condensates remain functionally competent, then SUMOylation may act as a molecular switch controlling PRC1 clustering and chromatin organization. Using RNAi-mediated depletion of SUMO in *Drosophila* wing discs, we examined how this global perturbation impacts PRC1 foci formation, chromatin topology, and gene expression. Our results reveal that SUMO-dependent regulation of PRC1 clustering is a key determinant of PcG-mediated chromatin organization, highlighting the profound architectural consequences of disrupting the SUMO pathway, which correlates with changes in gene expression within PcG target domains. These findings have potential implications for developmental gene regulation and the dynamic control of nuclear architecture during cell physiology and differentiation.

## Results

### Loss of SUMOylation transforms PRC1 condensate morphology and dynamics

To decipher the mechanism of Polycomb foci formation and address the role of PRC1 phase separation, *in vivo,* we took advantage of the previously reported observation that Pc-GFP accumulates in large, intense nuclear foci upon SUMO inactivation^15^. We compared the formation of Polycomb foci in fly lines that constitutively expressed Pc-GFP where we either induced a SUMO-KD or overexpressed Pc-GFP in the pouch of wing imaginal discs. Compared to control discs, Airy Scan imaging confirmed that SUMO-KD induces the formation of few, large (frequently a single one), intense Polycomb foci per cell nucleus. On the other hand, overexpression of Pc-GFP increases the intensity of all Polycomb foci (**Fig. 1a**) without merging them into fewer and larger ones. This result indicates that the formation of large Polycomb foci in SUMO-KD does not simply result from an increase in the nuclear concentration of Pc-GFP, but suggests that SUMO-KD specifically affects the condensation properties of Pc-GFP.

**Figure 1:**
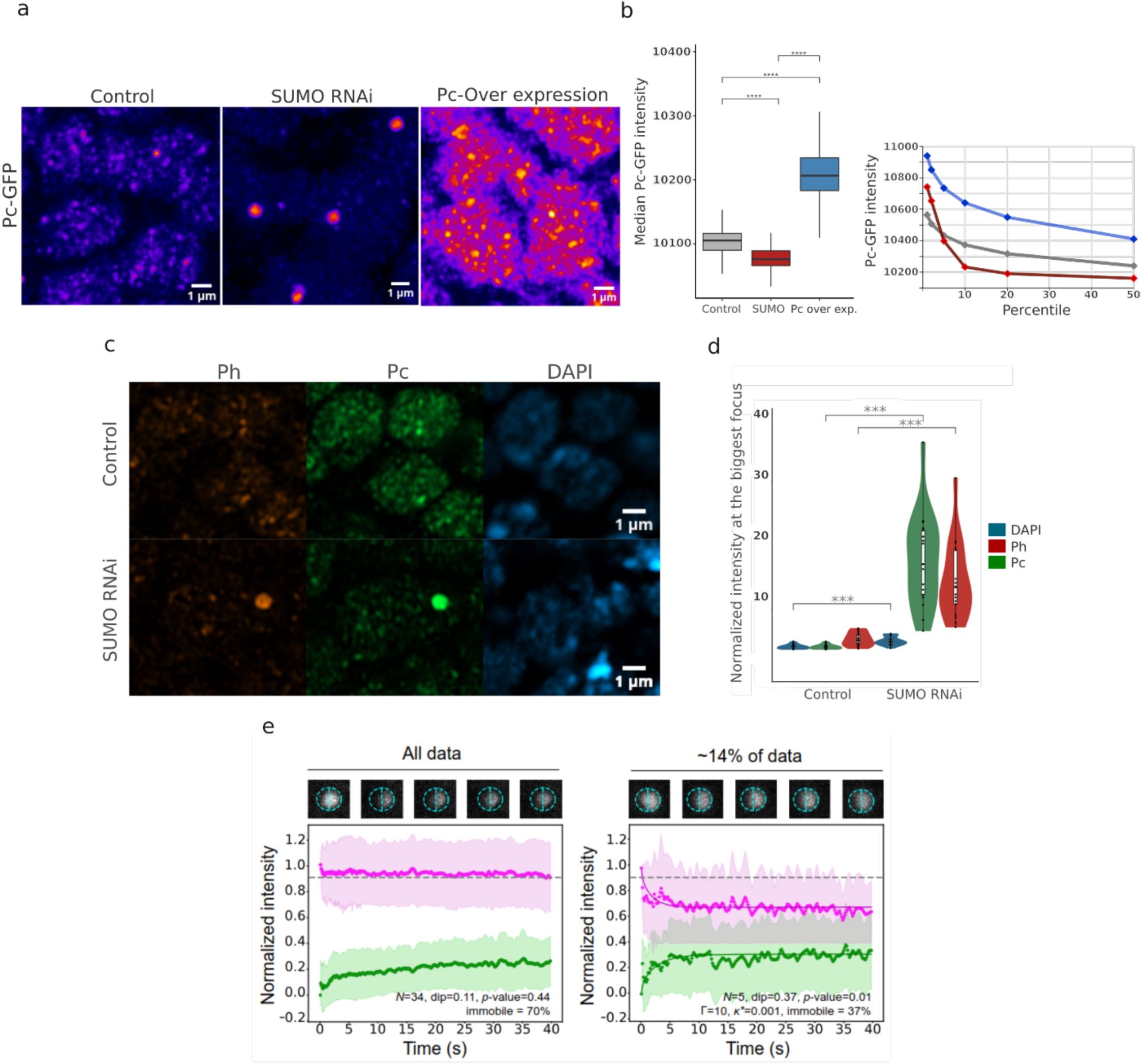
SUMO RNAi leads to unusually big PRC1 foci formation. **(a)** 2 µm thick projections showing the nuclear distribution of Pc-GFP in wing discs of control, SUMO-KD and overexpressing Pc-GFP flies. The pseudo-color images show that few Pc-GFP foci become very intense in the SUMO-KD, while the intensity of all foci increases when Pc-GFP is overexpressed. Scale bar: 1 µm. **(b)** Box plots comparing median Pc-GFP intensities in cell nuclei of control (grey), SUMO-KD (red) and overexpressing Pc-GFP (blue) wing discs (left). Compared to control wing discs, the intensity of nucleoplasmic Pc-GFP significantly decreases in SUMO-KD, whereas it strongly increases when Pc-GFP is overexpressed. Cumulative histograms comparing the intensity of Pc-GFP foci in control (grey), SUMO-KD (red), and Pc-GFP-overexpressing (blue) wing discs (right). Pc-GFP foci are ranked in descending order of intensity, and the points corresponding to the first, second, fifth, tenth, twentieth, and fiftieth percentiles are shown.**(c)** Airyscan images of Immuno-staining against Ph, Pc and DAPI staining from left to right, in control sample (top) and SUMO RNAi sample (bottom). **(d)** Quantification of the immunostaining in Control and SUMO RNAi samples Pc (green), Ph (red), DAPI (blue). Normalized intensity indicates the intensity of the signal within the largest focus in the nucleus, divided by the total intensity inside the nucleus for the respective signal. **(e)** Average MOCHA-FRAP curve of all the analyzed foci (n=34, dip=0.11, p.value=0.44). The dip depth with a *p*-value above 0.05 indicates that the majority of the PRC1 foci do not show liquid-like behavior with an interfacial barrier (left). MOCHA-FRAP curves of a subpopulation of 5 foci (that are included in the average curve shown on the left) that exhibit liquid-like behavior with an interfacial barrier (n=5, dip=0.37, p.value=0.01) (right). Solid lines represent the theoretical recovery for an internally mixed condensates with a Pc-GFP enrichment of Γ=10 and a vanishing permeability of *κ*\*=0.001. In both panels, the curve in purple indicates the change in intensity of the non-bleached half of the focus, and the curve in green indicates the recovery of the intensity of the bleached half of the focus. The fraction of immobile molecules in the focus can be detected by the final distance between the purple and the green curves, as indicated in the panel.

Since phase behavior is often concentration-dependent^19,20^, we asked whether SUMOylation affects Pc protein abundance. To estimate the free amount of nuclear Pc-GFP, *i.e.* the fraction of Pc-GFP not located in Polycomb foci, we measured the median intensity of Pc-GFP in the cell nuclei of control, SUMO-KD and Pc-GFP overexpressing wing discs. In addition to the formation of few large intense Polycomb foci, SUMO-KD also significantly reduced the level of Pc-GFP outside foci, compared to control conditions (**Fig. 1b**), suggesting that SUMO inactivation favors the association of Pc-GFP with foci. On the other hand, Pc-GFP overexpression increases the level of free Pc-GFP in cell nuclei (**Fig. 1b**). To quantify the accumulation of Pc-GFP in Polycomb foci, we computed local maxima to detect Pc-GFP foci and then ranked their intensity in descending order to compare the control, SUMO-KD and overexpression conditions. The intensity of Pc-GFP in all nuclear foci is higher than that observed in control wing discs when Pc-GFP is overexpressed. Conversely, only a few Pc-GFP foci (approximately 5%) show a higher intensity in SUMO-KD wing discs than in control discs. The remaining foci show a lower intensity in SUMO-KD, suggesting that most of the Pc-GFP accumulates in a few intense foci (**Fig. 1a-c**). Thus, SUMO loss alters condensate properties rather than simply stabilizing Pc protein. To verify that the localization of Pc-GFP reflects the nuclear distribution of PRC1, we performed immunolabelling experiments using the PRC1 subunit Ph and counterstained them with DAPI. Super-resolution Airyscan imaging of both control and SUMO-KD wing discs shows a strong co- localization between Ph and Pc subunit in both conditions, demonstrating that the large, intense Polycomb foci observed upon SUMO-KD correspond to PRC1 structures (**Fig. 1c**). In addition, DAPI signal also co-localizes with the large PRC1 foci upon SUMO-KD, confirming their chromatin-associated nature rather than off-chromatin aggregation (**Fig. 1c-d, S1b-c**).

PRC1 condensates have been proposed to form via liquid–liquid phase separation (LLPS)^2–7,12,13,21,22^. To specifically test this point, we next asked whether SUMOylation regulates Pc-GFP phase behavior by performing calibrated half-FRAP (MOCHA-FRAP)^23^. The purpose of this assay is to distinguish between liquid-like condensates with a significant interfacial barrier (characterized by a dip in the unbleached half curve due to intermixing) and condensates where exchange with the nucleoplasm dominates, reflecting assemblies formed by simple chromatin binding or by polymer-polymer phase separation. Analysis of PRC1 foci revealed that the large majority lacked the characteristic dip of liquid-like condensates (**Fig. 1e**), suggesting restricted internal dynamics and a shift toward a gel-like or solid-like state upon SUMO loss. Consistent with this interpretation, FRAP analysis revealed a mobile fraction of only ∼30% in PRC1 condensates (**Fig. 1e**), indicative of a stable scaffold with limited molecular exchange. However, ∼14% of foci did show a small dip, suggesting that a sub-population of large Pc-GFP foci do have liquid-like properties upon SUMO KD. These experiments could only be performed on the wing discs where SUMO RNAi is induced and not in control wing discs because Pc- GFP foci are too small to be half-bleached in this tissue. However, upon SUMO depletion PRC1 condensates transform into clearly larger, morphologically distinct structures with a lower internal dynamics than control^15^, without the surface tension that would be expected from a phase-separated liquid structure (**Fig. 1e**). The observation of liquid behavior in few cases suggests the possibility of a dynamic regulation of the properties of PRC1 foci during the accumulation of Pc-GFP upon SUMO KD.

### Changes in PRC1 self-interactions can explain the aberrant clustering behaviour of PRC1 upon SUMO-RNAi

To investigate how SUMO depletion can lead to the formation of large PRC1 condensates, we developed a generic biophysical framework coupling the Liquid-Liquid Phase Separation (LLPS) properties of PRC1 molecules with chromatin mechanics (see **Methods**). We considered a coarse grained polymer representation of *Drosophila* chromosome 3R at 1 kbp resolution (29 Mbp), featuring multiple H3K27me3 marked domains that can specifically interact with *N*_*P*_ diffusible self-attracting PRC1 molecules (**Fig.2a**). Using this framework, we systematically investigated multiple parameter sets to characterize the formation of PRC1 foci and its impact on chromosome organization, in control and perturbed conditions (**Fig.2b-c, Fig.S2**).

**Figure 2:**
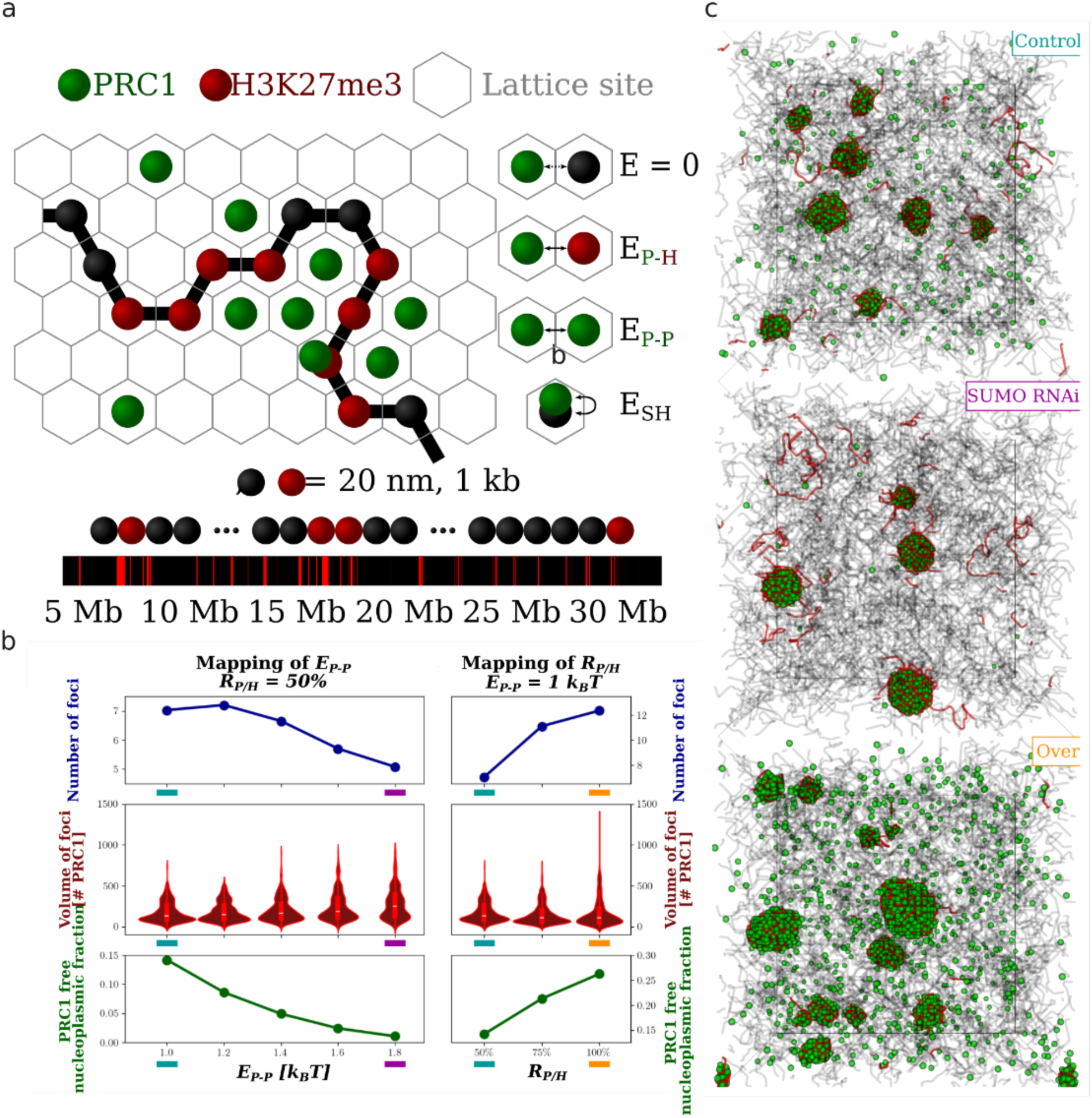
PRC1 self-interaction recapitulates the effect of SUMO RNAi. **(a)** Scheme of the chromatin and PRC1 simulation. Self-interacting PRC1 complexes, (in green, *E*_*P–P*_) exhibit interaction for H3K27-marked monomers (*E*_*P–H*_, in red). Steric hindrance (*E*_*SH*_) between PRC1 and monomer (both black and red) is taken into account. The 3R chromosome model is modeled from approximately 5 Mbp to 30 Mbp with methylation profiles approximating the size and location of PcG region in chromosome 3R of *Drosophila*. **(b)** Dependence of the number of foci, their volume distributions (in number of PRC1) and the PRC1 free nucleoplasmic fraction on the PRC1 energy of self-interaction *E*_*P–P*_ and on the ratio between PRC1 complexes and to H3K27 marked monomer *R*_*P*/*H*_. The ratio *R*_*P*/*H*_ is first fixed to 50% for the mapping of *E*_*P–P*_. The energy *E*_*P–P*_ is fixed to 1 k_B_T for the mapping of *R*_*P*/*H*_. The teal configuration is considered to be representative of the wild type situation. The purple and the orange configuration respectively are examples of the SUMO RNAi and the over-expressed configuration. **(c)** Example of the final frame for each configuration described in (b).

In particular, we tested the impact of the strengths of homotypic attractions between PRC1 complexes (*E*_*P–P*_) and of heterotypic interactions between PRC1 and H3K27me3-marked loci (*E*_*P–H*_) for different concentrations of PRC1 (given by the ratio *R*_*P*/*H*_ between *N*_*P*_ and the number of H3K27-marked monomers). As *E*_*P–P*_ increases (**Fig.2b**, left), the average number of foci decreases (top panel) with the appearance of larger foci at the expense of smaller ones (middle), while the free nucleoplasmic fraction of PRC1 is strongly reduced (bottom). As *E*_*P–H*_ decreases, foci are in average less numerous and larger and the free nucleoplasmic fraction of PRC1 increased (**Fig.S2**). All these effects are conserved regardless of the values of the other parameters (**Fig.S2**).

Therefore, the observations made by microscopy in SUMO RNAi conditions, *i.e.* the emergence of fewer but larger foci in a darker nucleoplasmic background (see **Fig 1**) are consistent with an increase of the strength of self-attraction *E*_*P–P*_ (**Fig.2c**). To further validate the model, we investigate the effect of an overexpression of PRC1 by varying *R*_*P*/*H*_(**Fig.2b,c**, right panels and **Fig.S2**). We predict an overall increase in the free nucleoplasmic fraction and in the number of condensates that become larger homogeneously, in perfect accordance to what was experimentally observed (**Fig. 1**). Taken together, polymer simulations indicate that homotypic interactions between PRC1s are necessary to explain the formation of large, intense PRC1 foci observed upon SUMO inactivation, providing indirect evidence that PRC1 phase separation also occurs *in vivo*.

In addition, all this suggests that SUMOylation may act as a solubility factor buffering the interactions between PRC1 molecules. Its removal leads to increased PRC1-PRC1 interactions, aberrant PRC1 clustering and the formation of large PRC1 foci.

### Loss of SUMOylation leads to changes in gene expression independent of H3K27me3

Given the dramatic reorganization of PRC1 condensates, we next asked whether SUMO depletion alters gene expression. We performed bulk RNA-seq on SUMO RNAi wing discs. Because SUMO is required for diverse cellular processes essential for viability^24–25^, we first confirmed that the tissues used were not undergoing widespread apoptosis. RNA levels of canonical apoptotic markers in *Drosophila*, including *rpr*, *Dronc*, *Dcp-1*, *AIF*, *Alg-2*, and *hid*, showed no significant upregulation at the developmental stage analyzed (**Fig. S3b**)^26,27^. Consistently, SUMO-depleted wing discs appeared overall healthy, although flies with SUMO RNAi either failed to hatch after the pupal stage or emerged with severe wing defects, suggesting that apoptosis might occur later in development. A modest increase in *rpr* expression is consistent with these tissues being primed for apoptosis while remaining healthy and viable at the time of analysis^26,27^. RNA-seq analysis further revealed extensive transcriptional changes, with 700 significantly differentially expressed (DE) genes, including known Polycomb targets such as *vg*, *Hr51* and *ftz* (**Fig. 3a**). We did not detect deregulation of the canonical PcG target gene *Ubx* in the wing discs, however, gene set enrichment analysis (GSEA) indicated a coordinated shift toward an active state of cellular growth and developmental reprogramming (**Fig. 3b** and **S3**), consistent with a deregulation of the PcG machinery ^28^.

**Figure 3:**
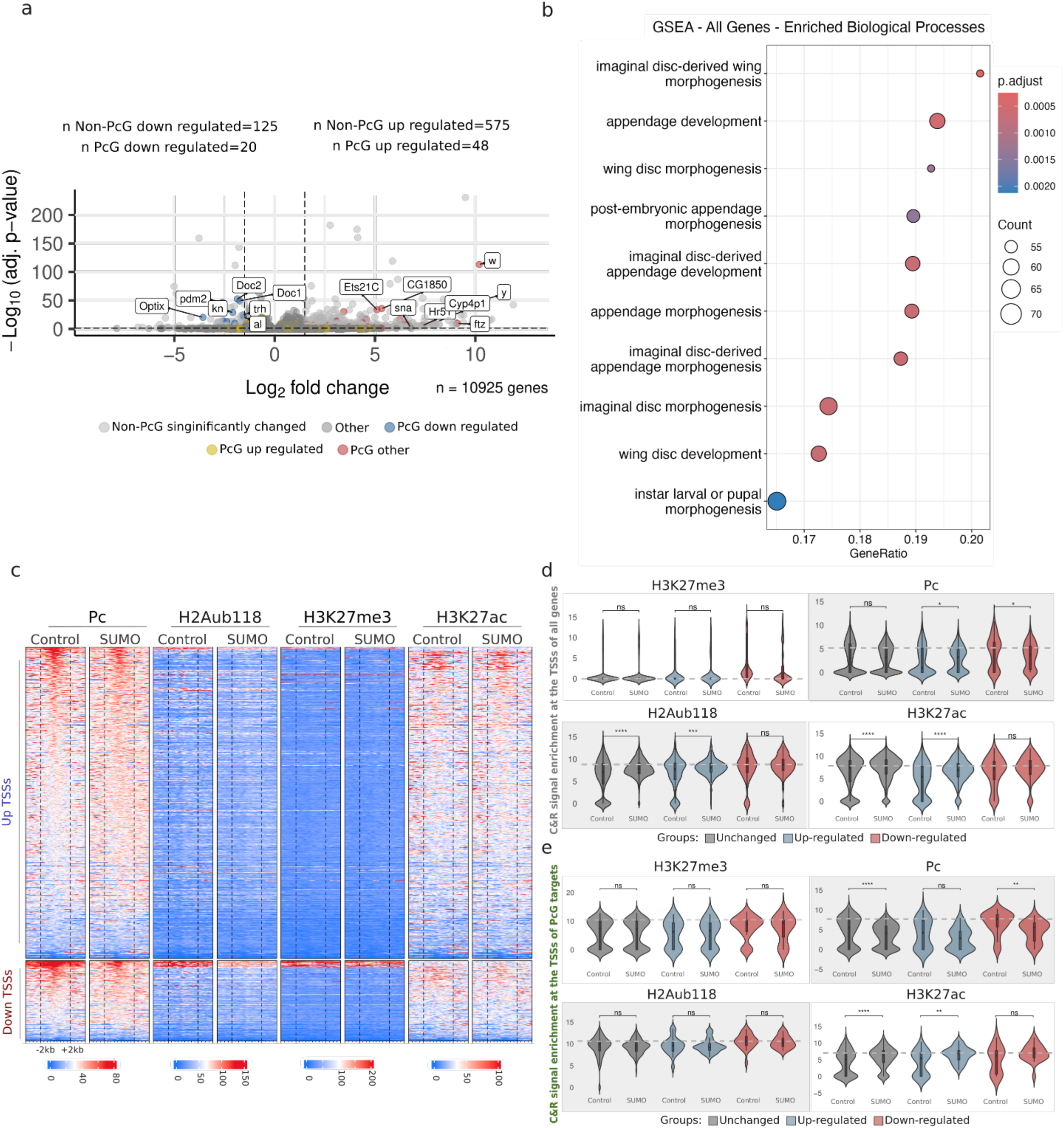
SUMO RNAi leads to gene expression changes without changes on H3K27me3 and H3K27ac marks. **(a)** Volcano plot of the differential gene expression (DE) analysis between SUMO RNAi and control sample; (p-adjust < 0.05 and absolute Fold-Change > 1.5). The PcG target genes are highlighted. **(b)** GSEA of all the DE genes. Top 10 enriched processes are represented. **(c)** Heatmaps of Pc, H2Aub118, H3K27me3 and H3K27ac marks around the TSSs of differentially expressed genes. Up TSSs= Transcription start site of up regulated genes; Down TSSs = Transcription start site of down regulated genes. **(d-e)** Comparisons of the normalized C&R signals around the TSSs of all (d) and PcG-target (e) genes between Control and SUMO RNAi. Grey indicates the TSSs of the genes which didn’t show a gene expression change; blue indicates the TSSs of the genes which were up-regulated; pink indicates the TSSs of the genes which were down-regulated. Dash lines indicate the median of the control sample.

We next asked whether these expression changes were associated with alterations in chromatin features. To address this point, we performed CUT&RUN profiling for Pc, H3K27me3, H2Aub118, and H3K27ac. Pc occupancy was mildly decreased across the genome and the Pc signal appeared slightly more diffusely distributed around TSSs in the SUMO-depleted condition (**Fig. 3c**). This reduction in Pc binding was observed even at PcG target genes that were transcriptionally downregulated (**Fig. 3e**).

On the other hand, the mild reduction in Pc occupancy was not accompanied by widespread changes in canonical Polycomb histone modifications. H3K27me3 enrichment was largely stable genome-wide, with only three novel sites appearing upon SUMO depletion (**Fig. 3c-d** and **Fig. S4**). PcG target genes also maintained their H3K27me3 signal (**Fig. 3e**). H2Aub118 showed a slight reduction around PcG target TSSs, but this trend was not statistically significant (**Fig. 3d-e**). Instead, H2Aub118 levels were increased near the TSSs of non-differentially expressed genes (**Fig. 3d**). As expected, H3K27ac levels were increased in SUMO RNAi compared to Control at TSSs of upregulated genes, but not at downregulated genes. However, we also detected a slight increase at unchanged genes (**Fig. 3c-e**) and importantly, no differences were found when we focused specifically on PcG target genes. Of note, the changes described above were modest, arguing against a major instructive role for histone modifications to drive changes in gene expression.

Together, these results indicate that the transcriptional changes triggered by SUMO depletion, especially down-regulation, do not follow canonical correlations with Polycomb histone marks. This suggested the hypothesis that altered PRC1 clustering may contribute changes in gene expression through mechanisms not directly dependent on H3K27me3 or H2Aub118 levels.

### Inter-TAD contacts are rewired upon loss of SUMOylation

The lack of major changes in H3K27me3 and H2Aub118, even at differentially expressed genes, suggested that the altered transcription upon SUMO depletion cannot be explained only by the altered levels of PRC1 or changes on SUMOylation of transcription factors. We hypothesized that SUMO RNAi-dependent changes in the nuclear organization of PRC1 might also alter the higher-order organization of PcG targets and contribute to gene expression changes. PcG domains are known to engage in long-range interactions between repressive TADs, often forming so-called “TAD cliques” that facilitate coordinated repression of target genes^29–30^. These PcG-associated TADs are typically enriched for H3K27me3 and depend on PRC1 for their long-range connectivity. We therefore asked whether SUMOylation contributes to 3D chromatin architecture.

To address this, we performed Hi-C experiments to obtain a genome-wide view of chromatin folding upon SUMO depletion (**Fig. 4a**). To systematically assess changes between the two conditions, we segmented the genome into physical domains (TADs). We classified them in different classes based on histone mark enrichment using ChromHMM (**Methods** and **Fig. 4a-c**): Active TADs (H3K27ac- enriched), Polycomb TADs (H3K27me3-enriched), constitutive heterochromatic TADs (H3K9me3- enriched), and null TADs lacking specific marks (**Fig. 4b**)^31^. We then quantified intra- and inter-TAD contacts in control and SUMO RNAi samples.

**Figure 4:**
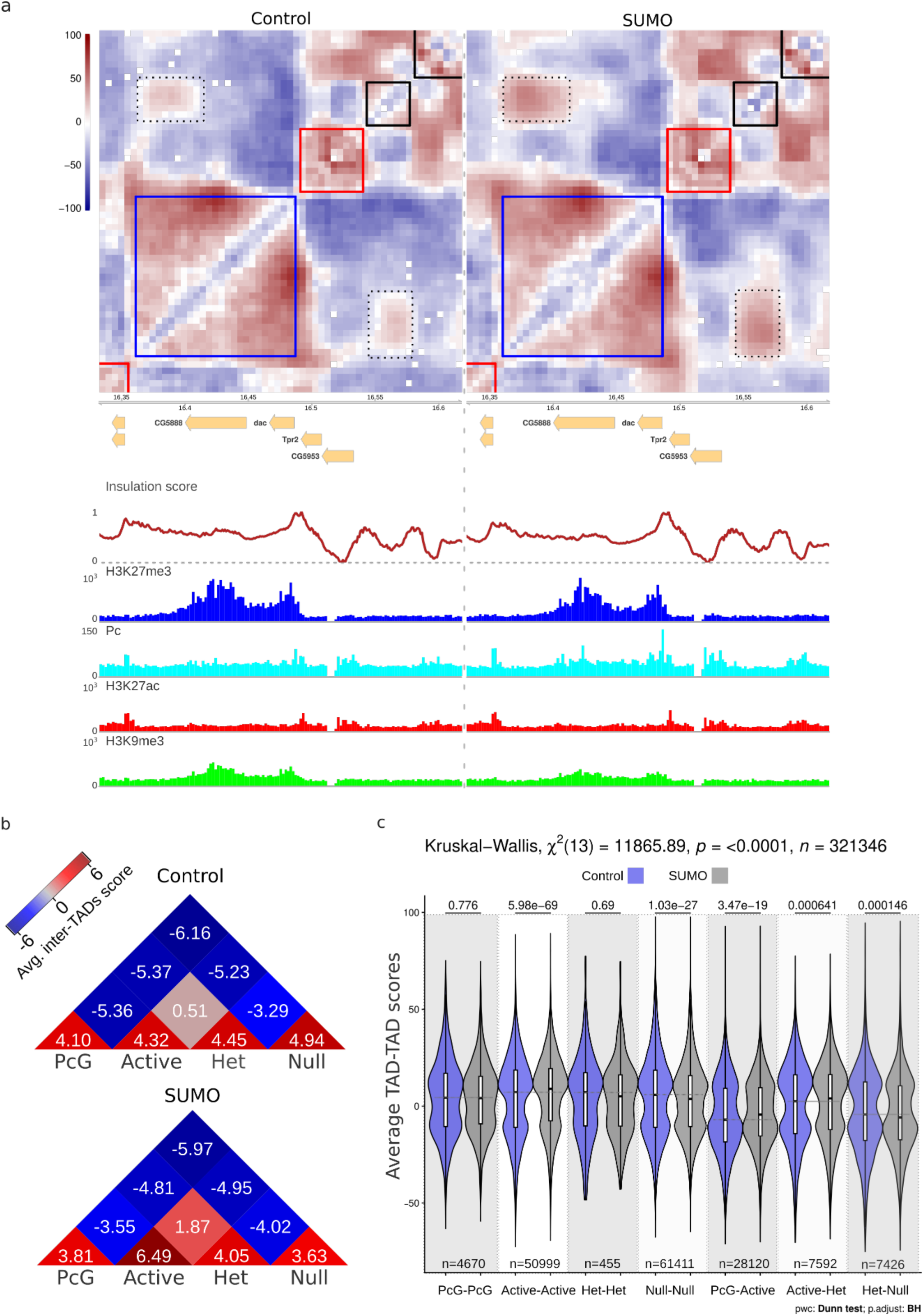
SUMO RNAi leads to genome-wide re-organization of the genome. **(a)** Heatmap of Hi-C scores in Control and SUMO RNAi samples for the genomic region around the *dac* locus. Colors towards red indicate high contact scores, colors towards dark blue indicate low contacts. C&R tracks for H3K27me3, Pc, H3K27ac, and H3K9me3 marks in control and the SUMO RNAi sample shown in the bottom. We used ChromHMM^27^ to assign chromatin states (colors) to each TAD in the genome (**Methods**). Chromatin states (colors) include: Active (red) chromatin enriched by H3K27ac, Null (black) chromatin not enriched in any features used for the analysis, HP1 (green) chromatin enriched by H3K9me3, and Polycomb (blue) chromatin enriched by H3K27me3. Each square indicates a TAD called and the color indicates the state. An example of increased TAD-TAD interaction between a PcG and Null TAD highlighted by the dashed lines. **(b)** Heatmap of the average inter-TAD score between the four chromatin states in Control and SUMO RNAi samples. **(c)** Quantification of the change of inter and intra-TAD contacts of each TAD type pair between control and SUMO samples. *P*-values are shown above each violin plot pair. *p*-values were corrected for multiple testing using Benjamini-Hochberg (BH) method.

This analysis revealed striking rewiring of inter-TAD interactions upon SUMO depletion. All TAD classes exhibited increased interactions with active chromatin, including Active–Heterochromatic and Active–Null contacts (**Fig. 4b-c**). In contrast, homotypic interactions of Null chromatin regions were reduced, while PcG-PcG and Het-Het (*e.g.*, Heterochromatin-Heterochromatin) contacts were preserved and, finally, Active–Active interactions became significantly more frequent. These results indicate that the effect of SUMO depletion extends beyond Polycomb chromatin and induces ectopic interactions with active chromatin, as observed between Active-PcG domains.

Thus, the deregulation of gene expression in SUMO-depleted wing discs in the absence of major changes to histone marks can be associated with the rewiring of TAD–TAD interactions. Concerning Polycomb chromatin, loss of SUMOylation may compromise the insulated chromatin environments normally maintained by PRC1 condensates, leading to exposure of PcG targets to neighboring active domains and a reconfigured regulatory landscape.

### SUMO KD leads to changes in PcG TAD interactions that correlate with PcG target gene expression

To further quantify and characterize the differential 3D chromatin interactions TADs, we analyzed contacts that each TAD makes with all other TADs.

We first focused our analysis on the PcG-PcG contacts. In order to analyze the relative changes in PcG chromatin contacts upon SUMO RNAi, we first divided PcG TAD pairs into quartiles based on the frequency of contacts made by the underlying PcG TADs. We then compared PcG TAD pair contacts between the control and the SUMO RNAi conditions, focusing on three categories (**Fig. 5a**): the entire distribution of inter-TAD contacts (All), the pairs that are weakly interacting, which are the pairs with little or no mutual contacts in controls (Q1 - 1st quartile), and the pairs that normally engage in strong interactions (Q4 - 4th quartile).

**Figure 5:**
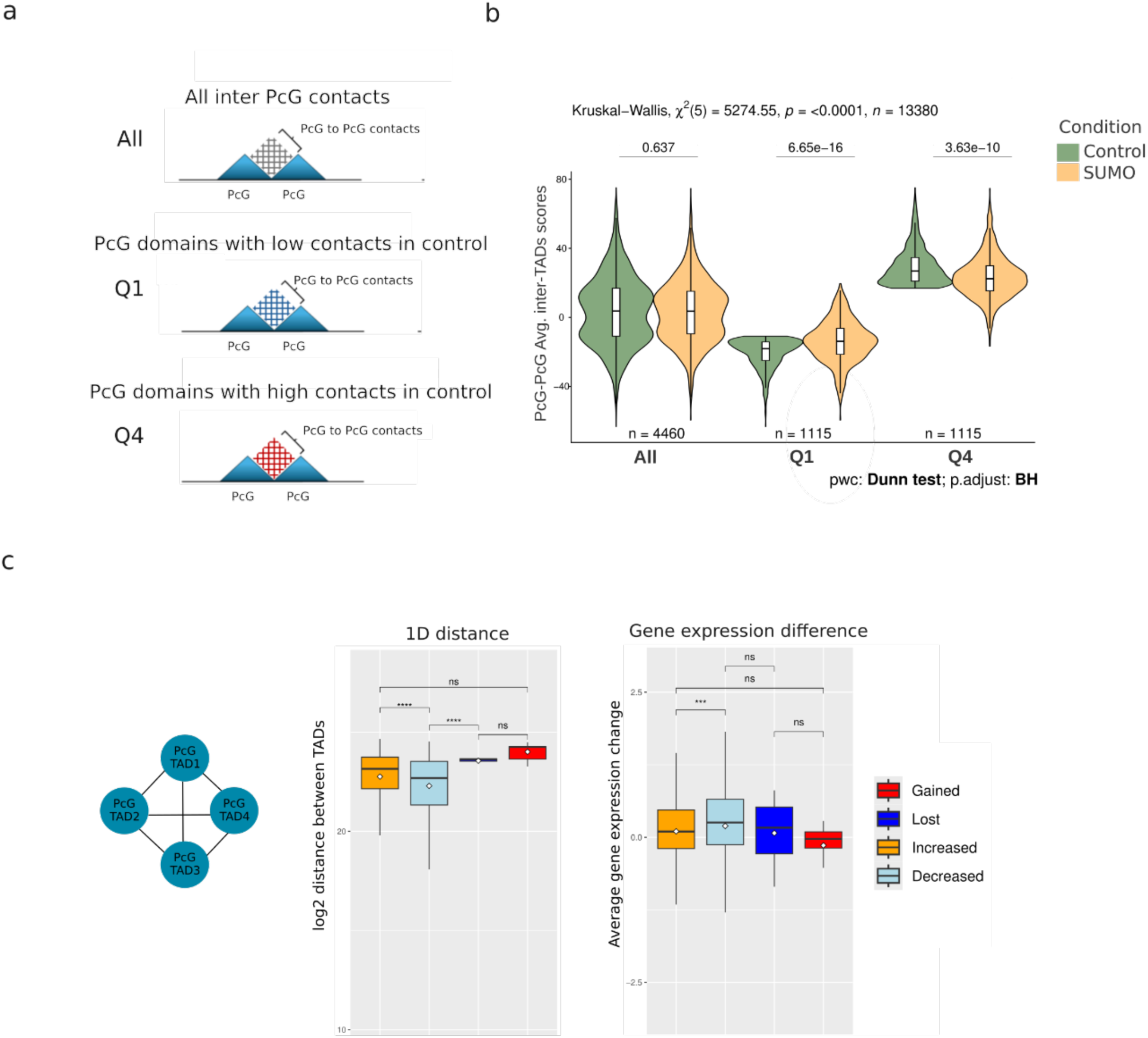
PcG-PcG TADs networks are re-wired upon SUMO RNAi. **(a)** Cartoon representing the stratification of inter-PcG-PcG TAD contacts. All indicates contacts without any filtering; Q1 represents the PcG TADs that engage in low contacts (below quartile 1) in the control sample; Q4 represents the PcG TADs that engage in high contacts (above quartile 4) in the control sample. **(b)** Quantification of the inter-PcG-PcG TAD contact scores (All, Q1 and Q4). **(c)** Comparison of different features of each TAD pair in different categories (increased, decreased, gained, lost). The left plot shows the log2 distance between each TAD, and the right plot shows the average gene expression change within each TAD. Average gene expression is calculated by the sum of log2 fold change of each gene in a particular TAD, divided by the number of the genes found in the respective TAD.

While the entire distribution of inter-PcG contacts showed no significant change, both the bottom and top quartiles displayed significant shifts (**Fig. 5b**). In particular, TAD pairs that had low numbers of PcG contacts in control tissues increased their contact frequencies upon SUMO RNAi, whereas TAD pairs with high PcG contact frequencies in controls decreased their contacts upon SUMO RNAi. This suggests that subsets of PcG domains alter their connectivity upon SUMO loss, potentially reshaping co-regulated gene groups. We then extended our analysis to heterotypic contacts between PcG TADs and other chromatin states (PcG–Active, PcG–Het, PcG–Null). PcG–Active contacts increased significantly (**Fig. S5**), consistent with the broader gain of interactions between Active domains and non-Active TADs observed in the genome-wide analysis (**Fig. 4b and c**). This reorganization likely perturbs the repressive environment normally required for PcG target gene silencing.

To further analyze the potential role of PcG-PcG contact changes on gene expression, we defined four categories to distinguish the differential the PcG inter-TADs contacts: gained (red), de novo contacts, which represents the TAD contacts which were not present in the Control but detected in the SUMO condition (ContactScoreControl = 0 & ContactScoreSUMO > 10); lost contacts which are not detectable in SUMO but detectable in Controls (ContactScoreControl > 10 & ContactScoreSUMO = 0), increased contacts which represent the TAD pairs which contact in both conditions but stronger in SUMO control (ContactScoreControl>0 & ContactScoreSUMO>0 & Δscore>10); and the decreased contacts which represent the weakening of the contacts that are present in both conditions (ContactScoreControl>0 & ContactScoreSUMO>0 & Δscore <−10). We show that TADs that gained contacts are further away from each other in linear (1D) distance1D genomic range, highlighting that loss of SUMOylation leads to longer-range contact increase, whereas the closer TADs decrease their contacts (**Fig. 5c**). Finally, the PcG TADs with decreased interactions in SUMO showed higher increase in the gene expression compared to the ones with increased contacts (**Fig. 5c**). In summary, these data indicate that SUMO depletion rewires PcG dependent genome contacts, affecting underlying gene expression patterns.

### PcG-Active TAD interaction changes correlate with PcG target gene expression

The significant overall increase in the PcG-Active contacts prompted us to investigate the contact patterns between these two types of TADs. First, we classified TAD pairs in quartiles based on the frequency of PcG-Active contacts made by each pair (**Fig. 6a**). The analysis of these data showed that, while there is a general increase in PcG-Active contacts when analysing all pairs, the increase is much stronger and more significant when restricting the analysis to the first quartile (Q1), the one with weakest contract frequency. In contrast, the analysis of Q4, the one with the highest contact frequency in controls, shows a significant reduction of contacts (**Fig. 6b**). These data suggest that the nuclear reorganization of PRC1 into large condensates induces a drastic remodeling of PcG TADs that disorganizes their overall contact pattern and exposes them to new subsets of contacts with active chromatin.

**Figure 6:**
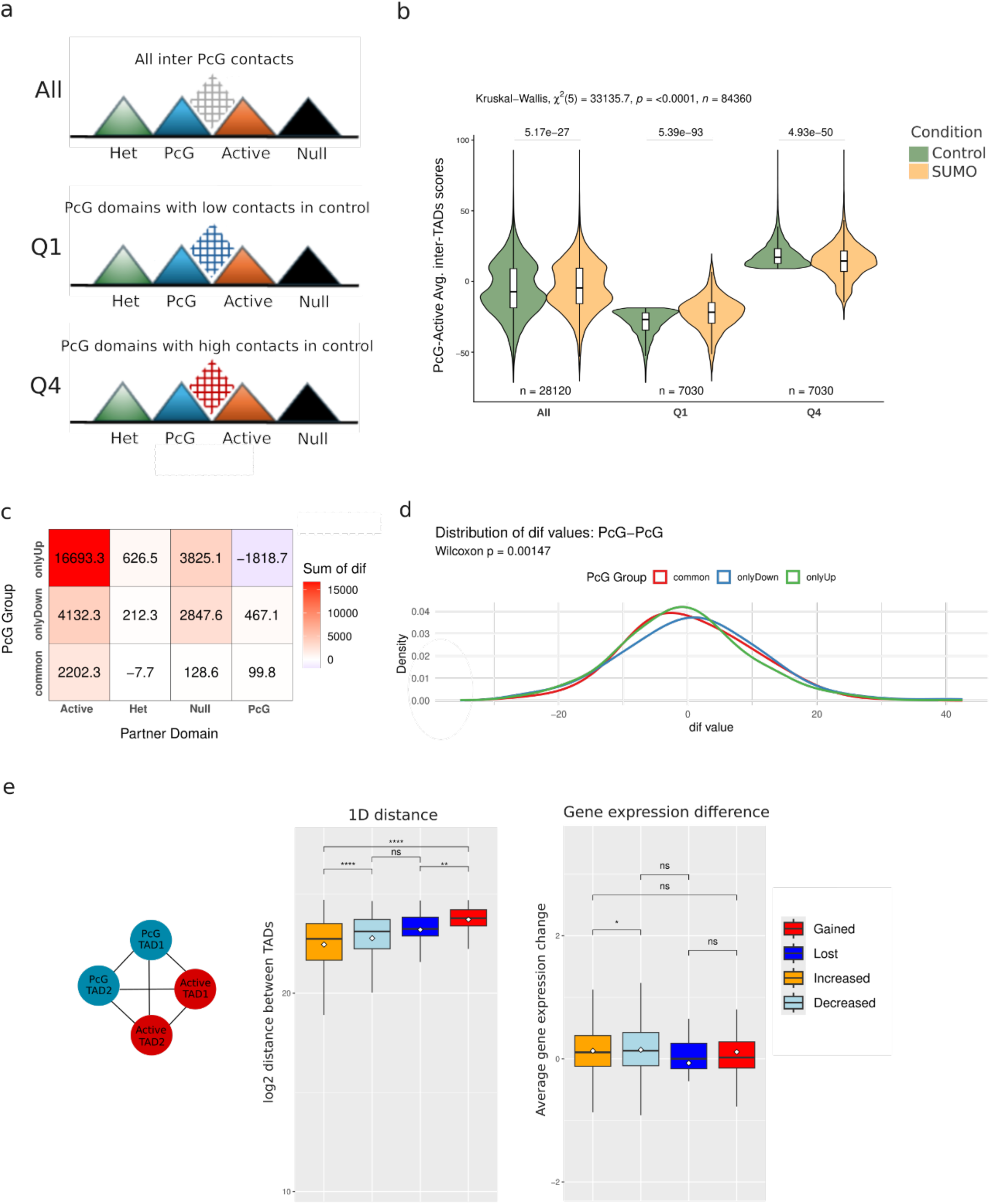
PcG-Active TADs networks are re-wired upon SUMO RNAi. **(a)** Cartoon representing the stratification of PcG-Active TAD contacts. All indicates contacts without any filtering; Q1 represents the PcG TADs that engage in low contacts (below quartile 1) in the control sample; Q4 represents the PcG TADs that engage in high contacts (above quartile 4) in the control sample. **(b)** Quantification of the contact scores between PcG and Active TADs in each category (All, Q1 and Q4). **(c)** Heatmap of the sum of the changes on each type of inter-TAD contacts. PcG TADs were split into three categories: those that contain only upregulated genes (onlyUp); only down regulated genes (onlyDown), or both down and up regulated genes (common). The total score changes for each of the three classes with all other TADs (Active, Het, Null, PcG) is indicated in each box in the plot. The score change is calculated for the TADs at the same chromosome arms only. **(d)** Distribution of the differences of contact scores of PcG-PcG TADs in each category. **(e)** Comparison of different features of each TAD pair in different categories (increased, decreased, gained, lost). The left plot shows the log2 distance between each TAD, and the right plot shows the average gene expression change within each TAD. Average gene expression is calculated by the sum of log2 fold change in each gene under a particular TAD, divided by the number of the genes.

To link structural changes in PcG TAD contacts with transcriptional outcomes, we stratified PcG TADs into three groups: those containing only down-regulated genes (onlyDown), those with only up- regulated genes (onlyUp), and those with a mixture of both kinds of genes. For each category, we compared SUMO RNAi and control samples by calculating the change in contact scores for every TAD pair and summing the differences by TAD class (e.g., PcG^onlyUp–active, PcG^onlyDown–null; **Fig. 6c**). All three PcG domain types exhibited increased contacts with active TADs, but the effect was most pronounced for onlyUp domains (**Fig. 6c**). By contrast, PcG–PcG interactions decreased in onlyUp domains but increased in onlyDown domains, a difference that was statistically significant (**Fig. 6c-d**). These findings suggest that SUMO depletion promotes exposure of PcG domains to active chromatin, while losing PcG-PcG interaction, inducing upregulation of genes in these PcG TADs.

Focusing on the relation between linear distances of TADs and changes in contact patterns, we found that Active–PcG TAD pairs with increased contact frequency are closer to each other on the linear scale along the chromosome than those with decreased contacts, indicating that heterotypic TAD interaction increases preferentially occur between nearby TADs (**Fig. 6e**). Furthermore, the average gene expression change was slightly lower in pairs of PcG-Active TADs that increase interaction, suggesting that an increase of Active TAD contacts with PcG TADs might reduce transcription, albeit marginally.

Finally, we analyzed whether SUMO depletion affects intra-TAD interactions within PcG domains. Pile- up plots and distributions revealed no significant changes in intra-PcG contact strength (**Fig. S6a** and **b**). Stratification by initial contact levels (low *vs.* high in control) similarly showed no differences (**Fig. S6b**). Thus, PcG domains maintain their internal insulation and contact patterns upon SUMO loss, showing that the main effect of PRC1 clustering upon SUMO depletion is to modify long-range chromatin interactions of Polycomb TADs rather than interfering with short-range chromatin fiber folding.

## Discussion

In this study, we demonstrate that global reduction of nuclear SUMOylation profoundly reorganizes the material properties of Polycomb Repressive Complex 1 (PRC1) condensates and their associated chromatin domains. Remarkably, these architectural changes and the resulting transcriptional misregulation occur without major alterations in PRC1 binding and H3K27me3 distribution, highlighting the importance of higher-order chromatin organization as a separable layer of Polycomb-mediated regulation.

### SUMOylation as a modulator of PRC1 condensate material state

PRC1 functions as a key architectural organizer, forming nuclear foci that facilitate long-range interactions between H3K27me3-enriched domains^1,6–9^. Our data show that SUMO depletion transforms these foci into enlarged, less mobile structures with properties consistent with a gel-like or solid-like state, exhibiting reduced molecular dynamics and limited internal mixing (**Fig. 1e**). In our biophysical model, we found that increasing the attraction between PRC1 complexes (E_P−P_), is key to mimicking a hypo-SUMOylated state, resulting in a transition from many small, local clusters to a few significantly larger foci (**Fig. 2b**). These large clusters could act as larger topological hubs, pulling more distant H3K27me3-marked regions closer into physical space (**Fig. 5b**). This could lead to a global increase in longer-range chromosomal contacts, especially among PcG-enriched domains and between PcG-enriched and Active domains (**Figs. 5-6**). This PRC1 reorganization correlates with widespread rewiring of TAD interactions, suggesting that the regulation of PRC1 condensate properties is essential for maintenance of the appropriate contact pattern between specific Polycomb domains (**Figs. 5-6**). SUMOylation provides a reversible, tunable mechanism to regulate condensate size and material properties. While previous work has emphasized protein-protein interactions among PRC1 subunits in foci formation^7,10,22,32^, our findings demonstrate that SUMOylation critically modulates condensate dynamics and that the formation of aberrant, gel-like condensates upon SUMO loss can drive ectopic 3D TAD interactions that compromise Polycomb domain insulation.

### SUMO gates proper 3D genome organization

We observed that PcG-enriched TADs normally engage in specific long-range interactions, consistent with previous reports^29,30^. Upon SUMO depletion, these networks are extensively rewired, with PcG TADs relatively close along the genomic sequence losing contacts with each other while gaining ectopic interactions with neighboring active chromatin (**Figs. 5-6**). This reorganization suggests that SUMOylation helps constrain PRC1-mediated interactions to appropriate chromatin partners in normal conditions, preventing promiscuous cross-talk between repressive and active compartments.

Our findings complement prior work showing that chemical disruption of PRC1 condensates abolishes contacts among PcG target regions^9^. Here, SUMO loss drives the opposite extreme, excessive clustering and rigidity, which similarly disrupts proper topology by creating aberrant bridges between normally segregated domains. This non-monotonic relationship highlights the fact that both the presence and the proper material state of PRC1 condensates are essential for correct genome folding.

### Mechanistic uncoupling from histone marks

A striking finding is the stability of H3K27me3 despite a slightly reduced Pc binding and 3D reorganization of TAD-to-TAD interactions. This demonstrates that H3K27me3 maintenance and higher-order chromatin interactions are mechanistically separable. The persistence of H3K27me3 in many domains, even as their spatial relationships change, suggests that this mark may be more stable than the condensates that organize it.

The global increase in H3K27ac upon SUMO depletion (**Fig. 3d**) suggests broader effects on chromatin-modifying activities, potentially through misregulation of other SUMOylated factors. However, the specific correlation between PcG-Active contact increases and upregulation of PcG target genes (**Fig. 6**) implicates architectural changes that are driven by aberrant PRC1 clustering as the dominant mechanism disrupting Polycomb-mediated repression in this context.

### Limitations and future directions

A key consideration is that we employed global SUMO depletion rather than PRC1-specific SUMO mutants. While this approach affects hundreds of SUMOylated proteins, the coherence of the phenotype-specific changes in PRC1 condensate morphology, coupled with specific rewiring of PcG-associated TAD interactions, suggests a direct functional link. Supporting this, overexpression of a Pc- 3KR mutant (lacking three SUMOylation sites) shows partially enlarged nuclear foci^15^. However, when we replaced the endogenous wt *Pc* gene with a Pc-3KR mutant form by CRISPR-dependent mutagenesis we failed to detect phenotypic effects as strong as in SUMO RNAi condition on imaginal wing discs or adult flies, suggesting that other SUMOylation sites in Pc and/or SUMOylation of other PRC1 subunits might complement for the reduction in Pc SUMOylation. Dissecting the specific SUMOylation sites within PRC1 remains an important aim for future research.

SUMOylation is dynamically regulated during the cell cycle and in response to stress^33–36^, suggesting that PRC1 condensate properties may be tuned in specific biological contexts. SUMO may also alter protein interaction networks by recruiting partners with SUMO-interacting motifs (SIMs); for example, CtBP localizes to Polycomb bodies in a SUMO-dependent manner^37,38^. Future profiling of the PRC1 interactome under SUMO-depleted conditions might reveal how PTMs reshape condensate composition and function.

### Conclusions and implications

Spatial organisation of chromatin together with its associated proteins, is strongly interlinked with their physical properties^39,40^. Our findings establish SUMOylation as a critical regulator of PRC1 condensate material properties, 3D genome architecture, and transcriptional fidelity. By demonstrating that aberrant condensate clustering leads to ectopic inter-TAD contacts and gene misregulation independently of canonical histone marks, we reveal a broader paradigm in which reversible PTMs fine-tune the material state and regulatory capacity of nuclear condensates. This work links phase-separation mechanics directly to chromatin topology and gene expression control, suggesting that post-translational regulation of biomolecular condensation represents a general strategy for coordinating nuclear architecture with transcriptional programs in metazoans. Future studies are needed to explore how fluctuating SUMO levels during development, cell cycle progression, and stress responses dynamically regulate nuclear condensates to maintain genome organization and function.

## Acknowledgments

We would like to thank the MRI and *Drosophila* facilities (BioCampus Montpellier), CNRS, INSERM and the University of Montpellier. We are grateful to the Genotoul bioinformatics platform Toulouse Occitanie (BioinfoGenotoul, https://doi.org/10.15454/1.5572369328961167E12) and the Pôle Scientifique de Modélisation Numérique (PSMN) of the ENS de Lyon for computational resources.

## Author Contributions

G.C., N.A. and T.C. conceptualized the study. N.A. performed the experiments. L.F and N.A. performed H2Aub118 CUT&RUN, and crosslinked Pc CUT&RUN. N.A. performed the remaining CUT&RUN experiments. P.S.P and D.J performed the biophysical modelling. M.D.S. performed the bioinformatics analysis of Hi-C, CUT&RUN, and RNA-seq experiments and N.A. performed downstream analysis of genomics experiments. F.E. and F.M. supervised MOCHA-FRAP experiments. T.C. performed the direct fixing and imaging of Pc-GFP expressing wing discs. N.A. wrote the manuscript. N.A., M.D.S., T.C. and G.C. edited the manuscript with inputs from all other authors.

## Methods

### Fly lines and husbandry

All *Drosophila* melanogaster stocks were maintained on standard cornmeal agar medium at 25 °C under a 12 h light/dark cycle. Gene knockdown and transgene expression were achieved using the GAL4/UAS system (Brand and Perrimon, 1994) with the nub-GAL4 driver obtained from the Bloomington *Drosophila* Stock Center (BDSC).

RNAi lines were sourced from the Vienna *Drosophila* RNAi Center (VDRC) and the Transgenic RNAi Project (TRiP; smt3 RNAi, stock JF02869). The following genotypes were used in this study:

- w[1118]; P{w[+mC]=UAS-Dcr-2.D}1; P{w[+mW.hs]=GawB}nubbin-AC-62 (nub-GAL4 driver line)
- PcGFP/CyO; Smt3RNAi/TM6B (SUMO RNAi)
- Actin5C-PcGFP/FM7 (overexpression of Pc-GFP)

Crosses were performed at 25 °C, and F1 progeny was used for downstream experiments. All fly lines used in this work are the same as the ones used in Gonzales et.al,2014.

### Immunostaining

Home made antibodies pre-cleaned prior to the staining procedure.For pre-cleaning of antibodies against PRC1 components, at least 20 larvae were dissected, retaining imaginal discs but removing fat tissue. Carcasses were transferred to 1.5-ml microcentrifuge tubes containing 750 µl PBS, and fixation was performed by adding 250 µl 16% paraformaldehyde (4% final) for 20 min at room temperature on a rotating wheel. Samples were washed three times with PBS, then permeabilized in 0.5% Triton X-100 in PBS (PBTr) for 1 h at room temperature with gentle rotation; the PBTr solution was replaced 2–3 times during this step. Tissues were subsequently blocked for 1 h in 3% BSA prepared in 0.1% PBTr at room temperature.

Primary antibodies were diluted 1:500 (Ph and Pc antibodies) in PBTr containing 1% BSA. Carcasses were incubated with the antibody solution for 2 h at room temperature, after which the antibody solutions were recovered and used for staining newly dissected samples.

For staining, newly dissected larvae were processed as above (4% paraformaldehyde fixation for 20 min, 0.5% Triton X-100 permeabilization for 1 h, and blocking in 3% BSA for 1 h) and then incubated overnight at 4 °C with the pre-cleaned antibody solutions. The following day, tissues were washed 4– 6 times with 0.1% PBTr at room temperature, and then incubated with secondary antibodies (1:200 dilution in PBTr containing 1% BSA) for 2 h at room temperature. Samples were washed at least three times for 15 min each in 0.1% PBTr.

Nuclei were counterstained with 1 µg ml⁻¹ DAPI in 0.1% PBTr for 20 min at room temperature, followed by three 15-min washes in 0.1% PBTr and three quick PBS washes without detergent. Wing discs were then dissected from carcasses in PBS and mounted in Vectashield (Vector Laboratories).

### RNA preparation for RNA-sequencing

Third instar larvae were dissected in Schneider medium. For SUMO RNAi and for respective control only male larvae were dissected. Total RNA was extracted using TRIzol reagent and extracted RNA were purified using RNA Clean & Concentrator kit (Zymo Research, #R1015). The sequencing was designed for mRNAs, thus, poly-A RNA enrichment, library preparation and illumina sequencing were performed by BGI. 6 GB of data was produced for each sample by BGI using paired end 150bp sequencing.

### RNA-seq analysis

RNA-seq samples were aligned using STAR aligner (v2.7)^41^. Subread (v2.0.6)^42^ was used to create the count table with the command *featureCounts -p --countReadPairs -s 2*. This count table served as the input for DESeq2 (v1.40.2)^45^. The volcano plots were obtained using the R library EnhancedVolcano (v1.20.0).

### CUT&RUN experiments

CUT&RUN experiments were performed as described by Kami Ahmad in protocols.io (10.17504/protocols.io.umfeu3n) with minor modifications. We dissected 20 Wing Discs for histone marks and 40 WDS for indirect DNA-binding proteins in Schneider medium centrifuged them for 3 min at 2000g and washed them twice with wash+ buffer before adding concanavalin A-coated beads. For experiments involving Pc, we included a 5-minute crosslinking step with 0.1% formaldehyde on the wheel at room temperature. MNase digestion (pAG-MNase Enzyme from Cell Signaling) was performed for 30 min at 4°C on shaker. After Proteinase K digestion, DNA was recovered using SPRIselect beads and eluted in 15 μl of Tris 10 mM. DNA libraries for sequencing were prepared using the NEBNext Ultra II DNA Library Prep Kit for Illumina. Sequencing (paired-end sequencing 150 bp, roughly 2 Gb per sample) was performed by Novogene (https://en.novogene.com/). The following antibodies were used: H3K27me3 (1:100, Active Motif, catalogue no. 39155), H2AK118Ub (1:100, Cell Signaling, catalogue no. 8240), H3K27ac (1:100, Active Motif, catalogue no. 39135), H3K9me3 (1:100, Active Motif, catalogue no. 39161), Pc (home made by G.Cavalli lab). All experiments were performed in biological duplicates.

### CUT&RUN analysis

CUT&RUN samples for H3K27me3, Pc, H2Aub118, H3K9me3, H3K27ac, and IgG in Control and SUMO RNAi imaginal discs were aligned using Bowtie2 (v2.4.2) with the options *-p 8 --local --very-sensitive-local --no-unal --no-mixed --no-discordant --phred33 -I 10 -X 700*. Low-quality reads were filtered with samtools (v1.9) using the command “samtools view -F 4 -h -q 30.” The resulting BAM files were sorted, indexed and de-duplicated using sambamba (v1.0.0) with commands *sambamba sort* and *sambamba markdup --remove-duplicates* respectively. To generate normalized BigWig files, the bamCoverage command from deepTools2 (v3.5.5) was used with parameters *--normalizeUsing RPKM --ignoreDuplicates*. *Macs3* (v3.0.0b1) was used to call peaks on each replicate with commands macs callpeak -t -f BAMPE -g 142573017 -q 0.01 -c ${controlBam} for Pc and H3K271c and macs3 callpeak -t ${sortedDedupBamPerReplicate} -c ${controlBam} -f BAMPE -g 142573017 -q 0.01 --broad --broad-cutoff 0.01 for H3K27me3, H2Aub118, and H3K9me3. Per each condition, the peaks per target were defined as the intersection between the peaks called in each of the replicates using the command *bedtools intersect* (v2.31.1). R library DiffBind (v3.10.1 on R v4.3.3) was used to find differential binding regions on the union of the peaks obtained for the SUMO RNAi and Control conditions. To define chromatin states in the Control condition, ChromHMM analysis was applied by first binarizing the signal in the replicate-merged .bam files of H3K27me3, Pc, H2Aub118, H3K9me3, and H3K27ac using IgG as a control track (command: *java -mx40G -jar ChromHMM.jar BinarizeBam -b 200 -p 0.0001 ./${bamFilesDir}/ cellmarkfiletable.tsv ./${binarizedBamFilesDir}*) and next by defining a 4-state model based on the binarized signal (command: *java -mx40G -jar ChromHMM.jar LearnModel -b 200 ./${binarizedBamFilesDir} ./${modelDir} 4 dm6*). The resulting four ChromHMM emissions were assigned to chromatin states: Active (red) chromatin enriched by H3K27ac, Heterochromatin (green) enriched by H3K9me3, Polycomb (blue) chromatin domains enriched by H3K27me3, and Null (black) chromatin not enriched in any features used for the analysis.

### Hi-C experiments

Hi-C was performed using the EpiTech Hi-C Kit (Qiagen) according to the manufacturer’s instructions with minor modifications.

Sixty wing discs were dissected in Schneider’s medium and transferred to 200 µl HBSS containing 10.8 µl 37% formaldehyde (2% final). Samples were homogenized using a Biomasher II (Funakoshi, 320103) and fixed for 20 min at room temperature. Crosslinking was quenched for 5 min with 100 µl 3 M Tris-Cl (pH 7.5), followed by addition of 650 µl HBSS and gentle inversion. The Collagenase treatment from the manufacturer’s protocol was omitted, as it reduced DNA yield. Samples were centrifuged (800 × g, 5 min, 4 °C), and 250 µl QIAseq Beads (equilibrated to room temperature) were added to the remaining ∼250 µl containing nuclei. After 10 min at room temperature, beads were magnetically separated and washed sequentially with 50 µl ice-cold PBS, 150 µl cold RNase-free water, and 50 µl cold Buffer C1. The mixture was incubated on ice for 10 min and then at room temperature for an additional 10 min before a final wash with 500 µl cold RNase-free water.

For chromatin digestion, beads were resuspended in 40 µl Hi-C Digestion Solution and incubated at 65 °C for 10 min, followed by cooling on ice. Subsequently, 4.4 µl 10% Triton X-100 and 4 µl Hi-C Digestion Enzyme were added, and samples were incubated at 37 °C for 2 h (600 rpm) and then at 65 °C for 20 min. Samples were placed on ice and could be frozen at this stage.

End labeling was performed by sequential addition of 6 µl Hi-C End Labeling Mix and 1 µl Hi-C End Labeling Enzyme, with gentle mixing and incubation at 37 °C for 30 min. For ligation, 350 µl Hi-C Ligation Solution was added, and samples were gently inverted and incubated at 16 °C for 2 h. Samples were cooled on ice before proceeding or frozen for later use.

For de-crosslinking, 10 µl Proteinase K was added, and samples were incubated at 56 °C for 30 min and at 80 °C for 90 min, then cooled to room temperature. DNA was precipitated by adding 40 µl 3 M sodium acetate (pH 5.2) and 280 µl isopropanol, vortexed briefly, and applied (including beads) to a MinElute® column (Qiagen). The column was washed with 0.75 ml Buffer PE, centrifuged (17,900 × g, 1 min), and eluted with 35 µl Buffer EB pre-warmed to 65 °C.

Recovered DNA was diluted to 130 µl and sonicated for 90 s using a Covaris S220 (140 W peak power, 10% duty factor, 200 cycles per burst). Sonicated DNA was purified by adding four volumes of Buffer SB1, loading onto a MinElute column, washing twice with 700 µl 80% ethanol, and eluting with 50 µl Buffer EB pre-warmed to 65 °C.

### Hi-C analysis methods

Raw data from Hi-C sequencing were processed using the ‘scHiC2’ pipeline. Valid interactions were stored in a database using the ‘misha’ R package (https://github.com/msauria/misha-package). Extracting the valid interactions from the *misha* database, the ‘shaman’ R package (https://bitbucket.org/tanaylab/shaman) was used for computing the Hi-C expected models, Hi-C scores with parameters *k* = 250 and *k_exp* = 500. Specifically, Hi-C scores quantify the contact enrichment (positive values) or depletion (negative values) of each bin of the map with respect to a statistical model used to evaluate the expected number of counts. To generate this expected model, we randomized the observed Hi-C contacts using a Markov chain Monte Carlo-like approach per chromosome. Shuffling was conducted such that the marginal coverage and decay of the number of observed contacts with the genomic distance were preserved but any features of genome organization (for example, TADs or loops) were not. These expected maps were generated for each biological replicate separately and contained twice the number of observed *cis* contacts. Next, the score for each contact in the observed contact matrix was calculated using the *k* nearest neighbors (kNN) strategy. In brief, the distributions of two-dimensional Euclidean distances between the observed contact and its nearest *k_exp* neighbors in the pooled observed and pooled expected (per cell type) data were compared, using Kolmogorov–Smirnov *D* statistics to visualize positive (higher density in observed data) and negative (lower density in observed data) enrichments. These *D* scores were then used for visualization (using a scale from −100 to +100) and are referred to as Hi-C scores in the text. Accordingly, the color scale of the Hi-C scores comprises both positive and negative values. When computing the differential Hi-C scores maps of, the reference dataset was used as the expected model. The box plots show the median (central line), the 75th and 25th percentiles (box limits) and 1.5 × IQR (whiskers). *.mcool* were obtained using *cooler*^44^ (v0.10.2) and *cooltools*^45^ (v0.7.0) and normalized via the Iterative Correction and Eigenvector decomposition algorithm (ICE) with default parameters (command “*cooler zoomify -r 5000 file.cool -o file.mcool --balance*”). The physical domains (TADs) were called in the Control condition using the TopDom algorithm (v0.10.1)^46^ as in the HiCExperiment R-package (v1.4.0)^47^ using a window size parameter of 5 on the *.mcool* files at 5 kilo-bases (kb) resolution. Each of the 5-kb TAD borders were then refined at 1 kb by looking at the 1kb point of maximum insulation.

### Image acquisition

For Confocal and the half-FRAP experiments LSM980 microscope was used. For half-FRAP, time series were recorded every 150,200,300,350 or 500 ms. Bleaching performed at the 3rd time point. The pixel size was 30nm for the images. The pin hole was increased to 150µm. AiryScan microscopy was performed using a LSM 980 microscope (Carl Zeiss Microscopy, Iena, Germany) with a AiryScan2 detector and a 63× PlanApo objective having a numerical aperture (N.A.) 1.4. Pc-GFP was excited with a 488 nm laser diode and we used the mode AiryScan SR to collect images with an optimal pixel size of 42 nm in x, y and 170 nm in z. Processing of raw images to produce AiryScan images was done with the Zen software controlling the microscope with a wiener filter automatically adjusted by the software.

### Image analysis

To characterize Pc foci inside cell nuclei (**Fig. 1**), we quantified 3D images acquired in Airy Scan microscopy by using the software Imaris 9.8.0 (Oxford Instruments, UK). We applied the spots option of the Imaris software to identify nuclear foci and measure their Intensity. To estimate the intensity of Pc-GFP inside the nucleoplasm, we used the median intensity of Pc-GFP inside cell nuclei.

### Polymer model

PRC1 complexes and chromatin are modeled following the framework developed in previous work^48^. Briefly, PRC1s molecules are represented as a lattice gas with *N*_*P*_ particles, *i.e*. as beads of effective size *20 nm* residing on the vertices of a face-centered cubic lattice^49^. Similarly, chromatin is described as a semi-flexible, self-avoiding polymer chain composed of *N*_*C*_ monomers evolving on the same lattice as the PRC1s. Each monomer encompasses *1 kbp* of chromatin (5 nucleosomes) and can be in one of two epigenetic states: H3K27me3-marked (Polycomb-regulated regions) or H3K27me3-unmarked (the rest of the chromatin). The Hamiltonian H of the system describing its dynamics is composed of four terms: *H* = *H*_*bend*_ + *H*_*P–P*_ + *H*_*P–H*_ + *H*_*SH*_.

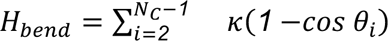 accounts for the bending rigidity of the chromatin where *θ*_*i*_ is the angle between monomers (*i* − *1*, *i*, *i* + *1*) and *κ* = *3.2 k*_*B*_*T* is the bending modulus of the polymer chain, corresponding to a Kuhn length of *100 nm*^50,51^.

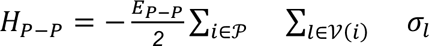 accounts for the self-attraction between PRC1 molecules where *E*_*P–P*_ is the strength of self-attraction, 𝒫 is the ensemble of PRC1s, 𝒱(*i*) are the lattice sites that are nearest-neighbor of the one where PRC1 *i* is seated, and *σ*_*l*_ = *1* if a PRC1 is on lattice site *l* (*σ*_*l*_ = *0* otherwise).

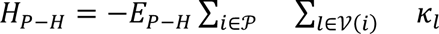 accounts for the cross-interaction between PRC1 molecules and H3K27-marked monomers where *E*_*P–H*_ is the strength of cross-interaction and *κ*_*l*_ = *1* if an H3K27- marked monomer is on lattice site *l* (*κ*_*l*_ = *0* otherwise).

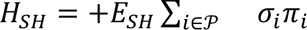 accounts for steric hindrance interactions between PRC1 molecules and any monomers where *E*_*SH*_ is the strength of steric hindrance and *π*_*i*_ is the number of monomer on the lattice site *i*.

The lattice is made of *N* vertices, with periodic boundary conditions. Each vertex has 12 nearest neighbors and can contain at most one PRC1 molecule and 2 chromatin monomers. Note that double occupancy by monomers is allowed if and only if monomers are consecutive along the chain, accounting for bond fluctuations while ensuring correct self-avoidance^48^.

### Monte-Carlo simulation

Simulations were performed starting from random, unknotted initial configurations for the chromatin chain and a uniform random distribution for PRC1 molecules. The system is evolved via a kinetic Monte-Carlo scheme, as detailed in previous work^48^. Briefly, each Monte Carlo step (MCS) consists of *N*_*C*_ monomer trial moves and *N*_*P*_ PRC1 trial moves. During a trial move, a PRC1 molecule or a monomer is randomly selected and an attempt to move it to one of randomly-chosen nearest neighbor vertex is attempted. The new configuration is accepted via the Metropolis criterion based on the energy difference between the trial and current configurations.

For each investigated parameter set, we simulated *N*_*traj*_ = *40* stochastic trajectories, each composed of a first warm up of *10⁶ MCS* in absence of self- and cross-interactions to relax the system, followed by *10⁷ MCS* with the full Hamiltonian, where snapshots were extracted every *10⁵ MCS*.

We simulated chromosome arm 3R (*N*_*C*_ = *29184* monomers) including *2859* H3K27-marked monomers distributed in several domains across the polymer based on ChIP-seq data in the wing disk. The ratio *R*_*P*/*H*_ between *N*_*P*_ and the number of H3K27-marked monomer was varied from *50%* to *100%*, *E*_*P–P*_ from *0.2* to *2.2 k*_*B*_*T*, *E*_*P–H*_ from *0.8* to *1.2 k*_*B*_*T* and *E*_*SH*_ from *8* to *10 k*_*B*_*T*. The set of parameters (*E*_*P–P*_ = *1 k*_*B*_*T*, *E*_*P–H*_ = *0.8 k*_*B*_*T*, *E*_*SH*_ = *8 k*_*B*_*T* and *R*_*P*/*H*_ = *50%*) is representative of the wild-type condition, capturing the strength of typical interactions of architectural proteins having LLPS properties^2,48,52^ and typical in vivo concentration^53^. Our results are not qualitatively dependent on this choice.

### Data analysis of the simulations

For each snapshot, PRC1 foci were defined similarly as in previous work^48^. Briefly, for each PRC1 molecule *i*, we defined its local PRC1 density 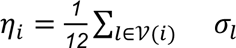 and its local H3K27-marked monomer occupancy ratio 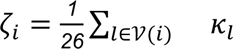. The ensemble of PRC1s belonging to a condensate is defined as 𝒟 = {*η_i_* + 𝜁_*i*_ ≥ 0.5, *i* ∈ 𝒫}.

The connected components of 𝒟 are the foci. The volume of each focus is defined as the number of PRC1s inside it. The PRC1 free nucleoplasmic fraction is computed as (*N*_*P*_ − *Card*(𝒟))/*N*_*P*_. The PRC1 free nucleoplasmic fraction, the number of foci and the distribution of foci volume were averaged over the 10 last frames of each simulation.

## Supplementary Figures

**Fig. S1.**
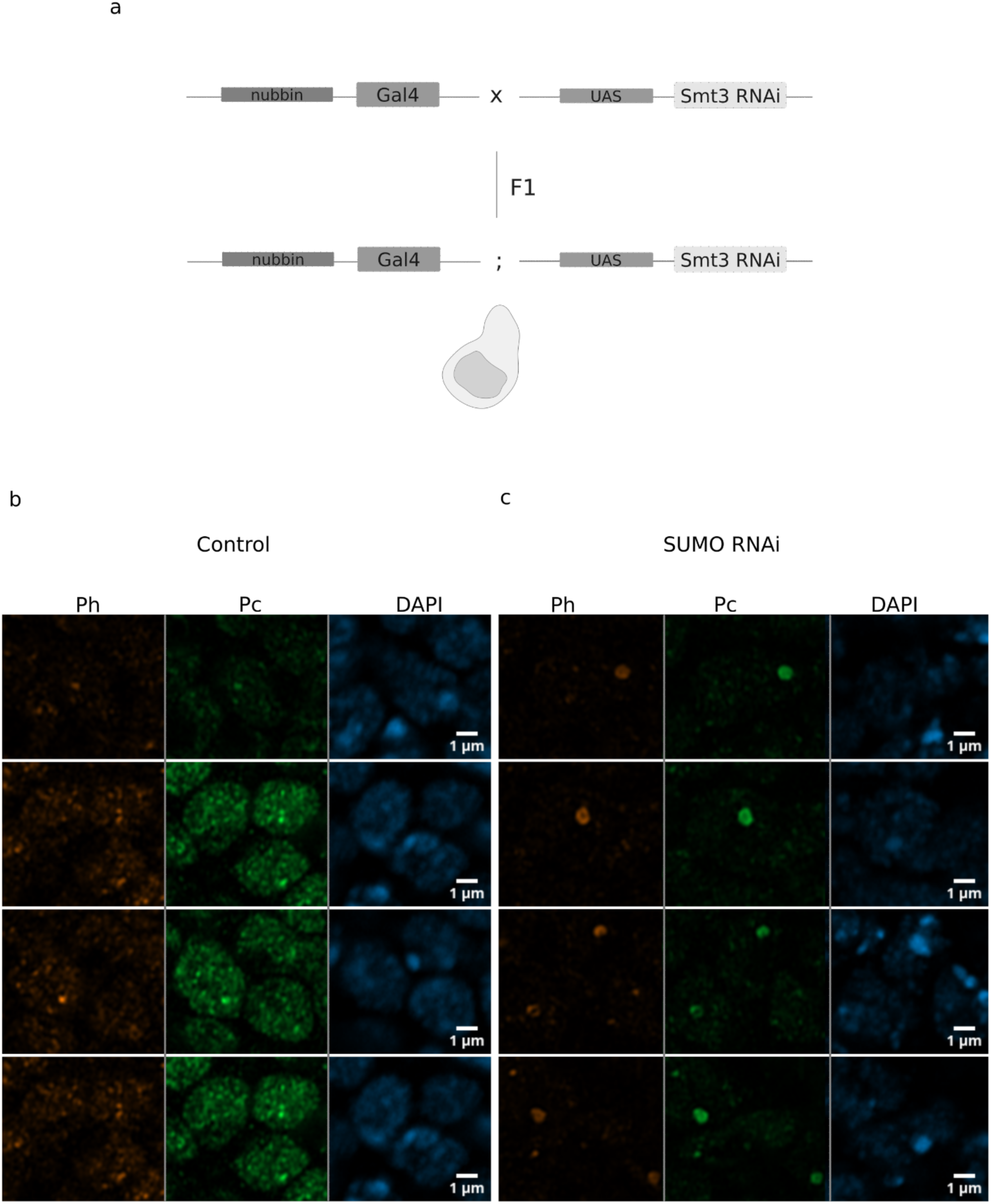
SUMO RNAi induced on the imaginal wing disc pouch leads to reorganization of PRC1 foci. **(a)** Cartoon representing the fly lines used to induce SUMO RNAi on *Drosophila* wing discs. **(b)** Airyscan images of Immuno-staining against Ph, Pc and DAPI staining, in control wing discs. **(c)** Airyscan images of Immuno- staining against Ph, Pc and DAPI staining, in SUMO RNAi wing discs.

**Fig. S2.**
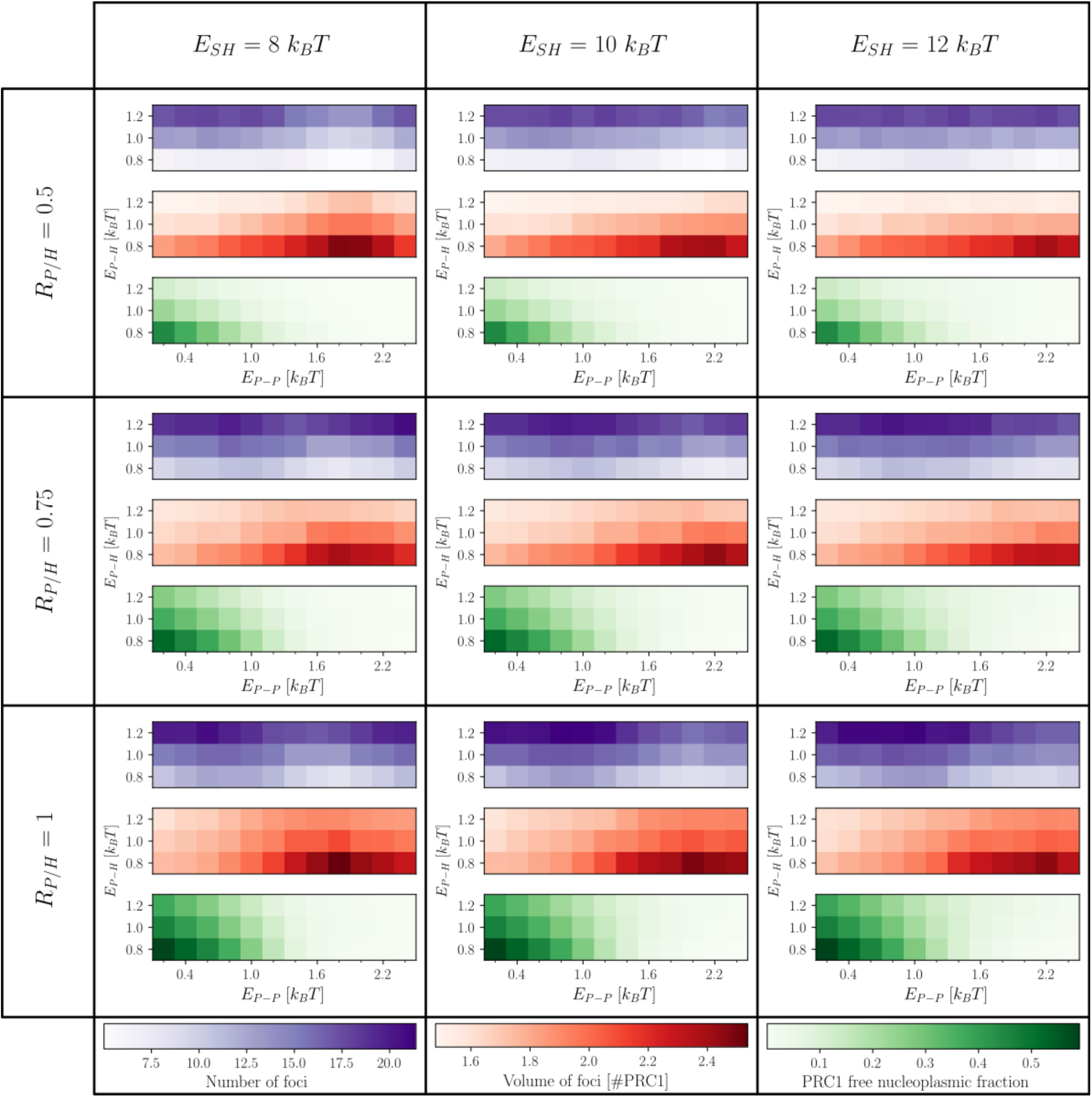
Table of simulation results of the biophysical framework. (Columns) Each column shows simulations of varying strength of steric hindrance (*E*_*SH*_). **(Rows)** Each row shows simulations of varying ratio between PRC1 complexes and to H3K27 marked monomer (*R*_*P*/*H*_). **(Heatmaps)** Each heatmap shows simulations of varying strength of PRC1 complexes self-interaction (*E*_*P*–*P*_, horizontal) and of varying PRC1 interaction strength for H3K27-marked monomers (*E*_*P*–*H*_, vertical). **(Blue)** Each blue heatmap shows the number of foci at the end of the simulations. **(Red)** Each red heatmap shows the average volume of foci at the end of the simulations in number for PRC1 complexes. **(Green)** Each green heatmap shows the average PRC1 free nucleoplasmic fraction. The trends shown in **Fig.2b** are not dependant on the values of *E*_*SH*_ and *R*_*P*/*H*_. Indeed, an increase of *E*_*P*–*P*_ leads to an in, the number of foci and the PRC1 free nucleoplasmic fraction are decreasing whereas the volume of foci is increasing.

**Fig. S3.**
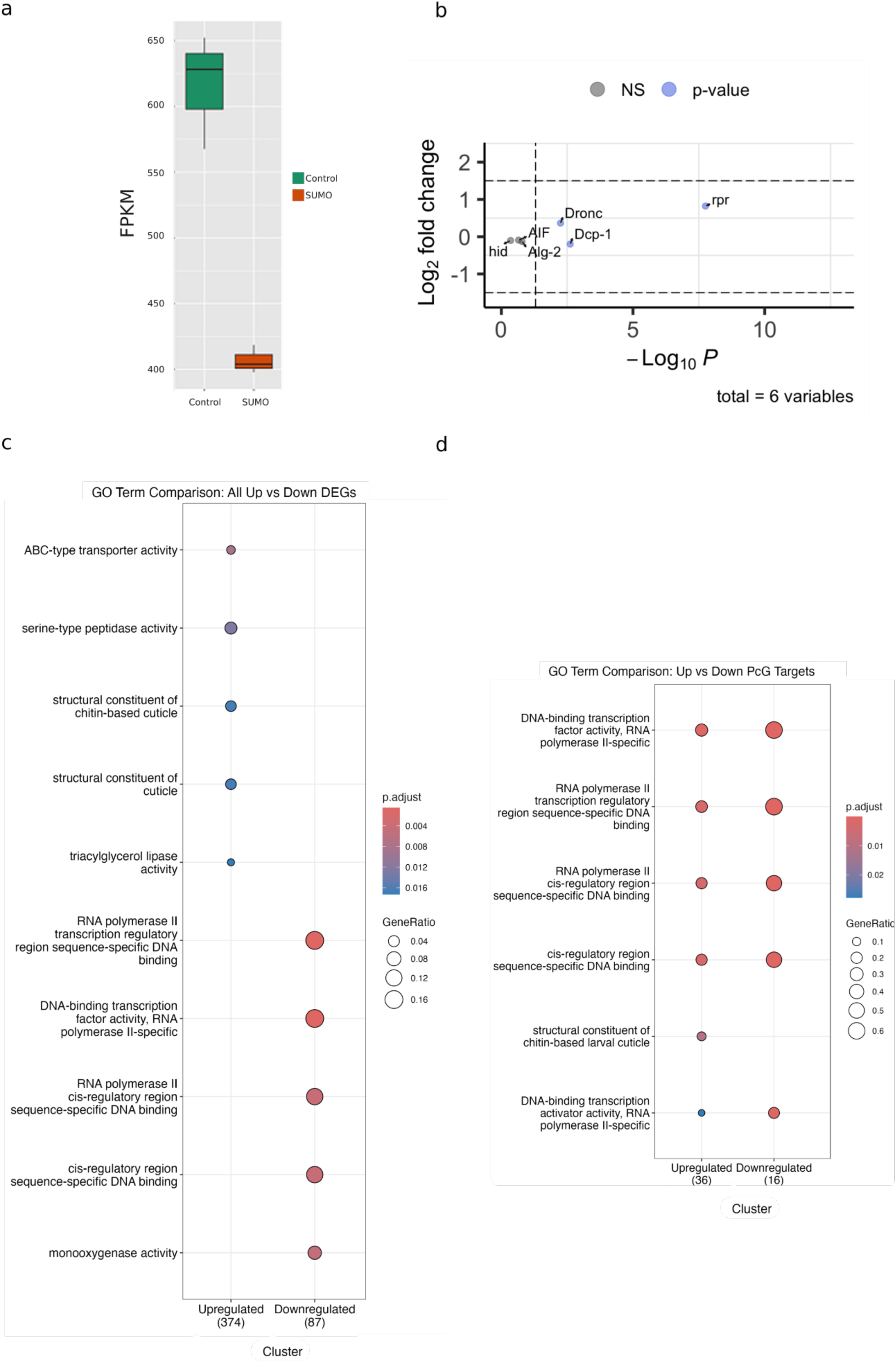
Gene expression profiles upon SUMO RNAi. **(a)** FPKM values of the SUMO gene (*smt3*) in the control and SUMO RNAi samples. P.adjusted= 6.49e-22. **(b)** Volcano plot showing the DEseq2 results for the apoptotic markers. Absolute value of the Fold change > 1.5, p.adjusted < 0.05. GO term enrichment analysis for the all differentially expressed genes upon SUMO RNAi. **(d)** GO term enrichment analysis for the differentially expressed PcG target genes upon SUMO RNAi.

**Fig. S4.**
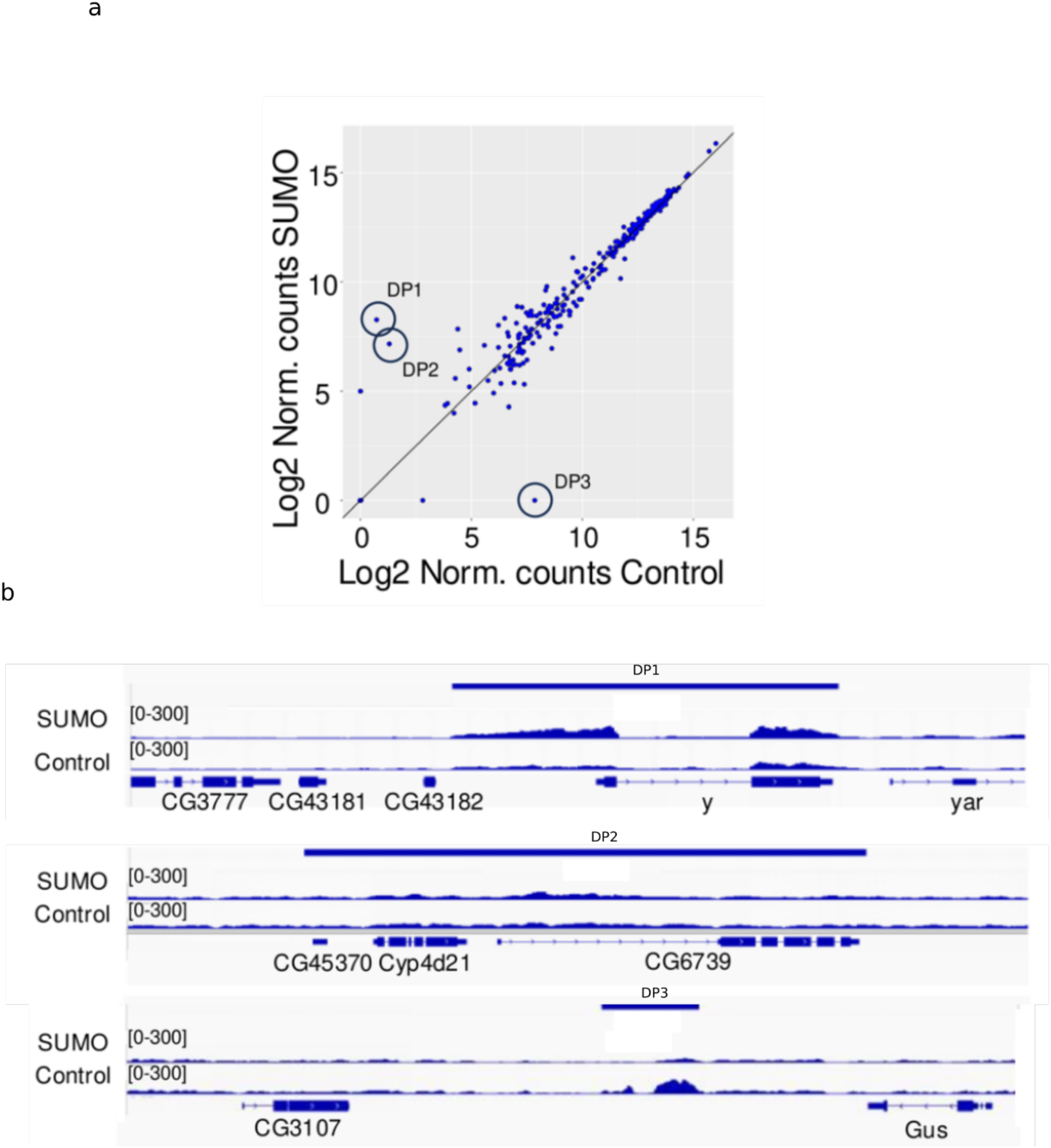
SUMO RNAi does not lead to major changes on the distribution of the H3K27me3 mark compared to the control. **(a)** Scatter plot of the enrichment of H3K27me3 mark, x axis indicates SUMO RNAi and y axis indicates the control. **(b)** Tracks of the differentially bound sites for the H3K27me3 mark in each condition.

**Fig. S5.**
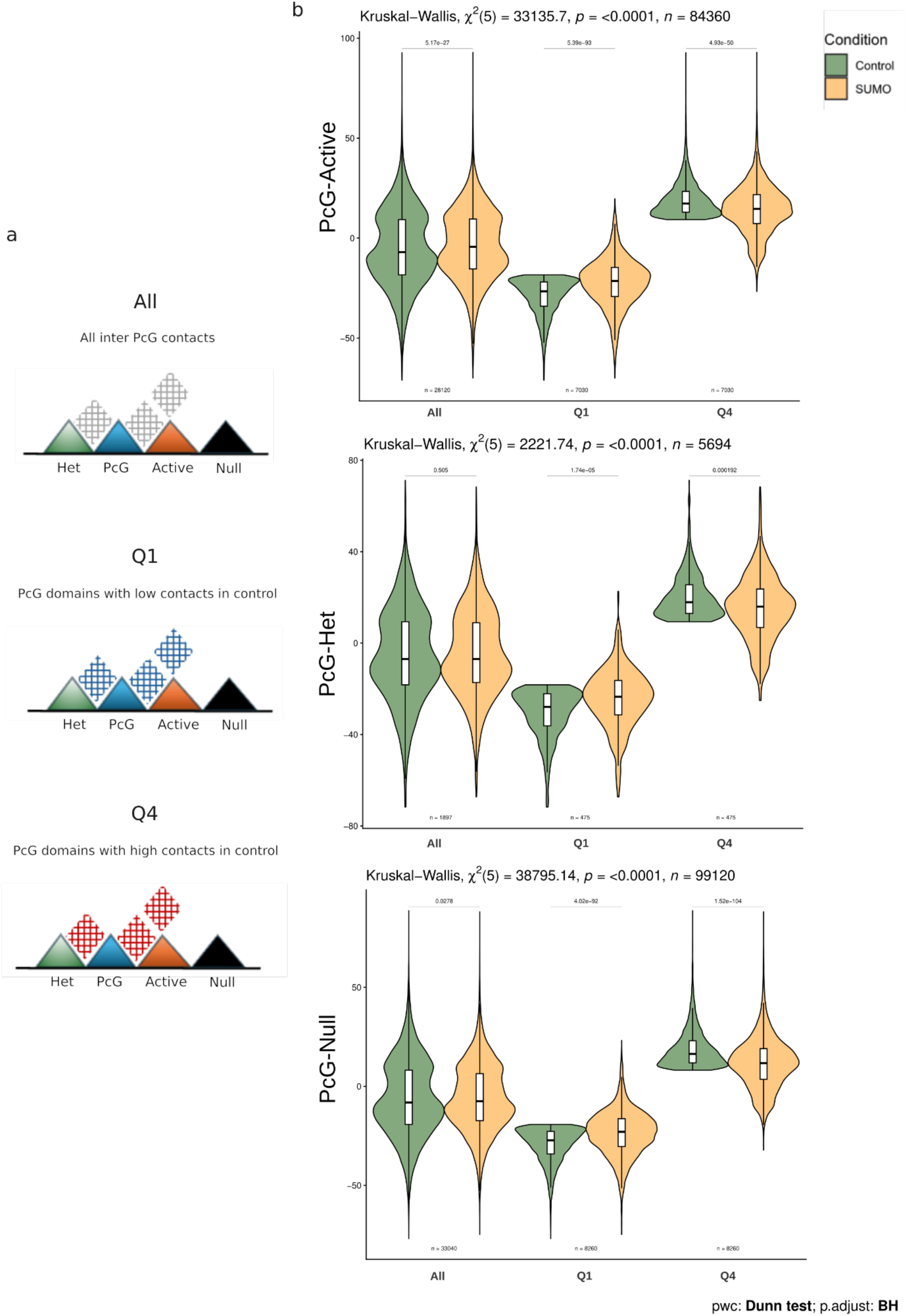
Heterotypic inter-PcG TADs networks are re-wired upon SUMO RNAi. **(a)** Cartoon representing the stratification of inter-PcG TAD contacts. All indicates contacts without any filtering; Q1 represents the PcG TADs that engage in low contacts (below quartile 1) in the control sample; Q4 represents the PcG TADs that engage in high contacts (above quartile 4) in the control sample(left). **(b)**Quantification of the contact scores between PcG TADs with other types of TADs in each category (all;Q1 and Q4).

**Fig. S6.**
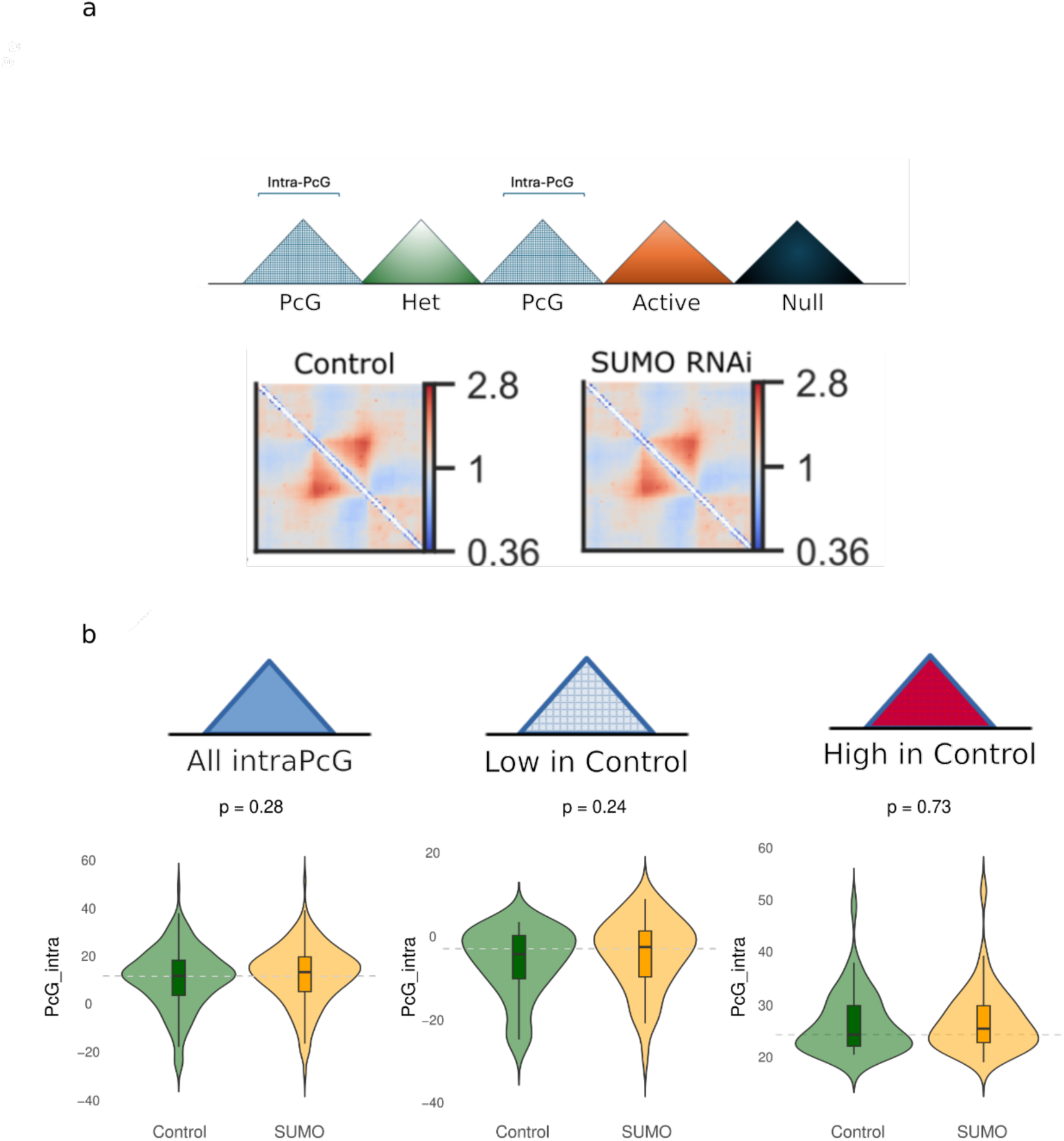
Intra-domain interactions of PcG domains don’t show significant changes upon SUMO RNAi. **(a)** Cartoon representing the intra-domain contacts that are measured (top); pile-up plots of PcG TADs in control and SUMO RNAi samples (bottom). **(b)** Cartoon representing the stratified intra-domain contacts (all intra-PcG TADs; Q1 intra-PcG TADs; Q4 intra-PcG TADs) that are measured (top).; quantification of the stratified intra TAD contacts (all intra-PcG TADs; Q1 intra-PcG TADs; Q4 intra-PcG TADs) (bottom).

**Fig. S7.**
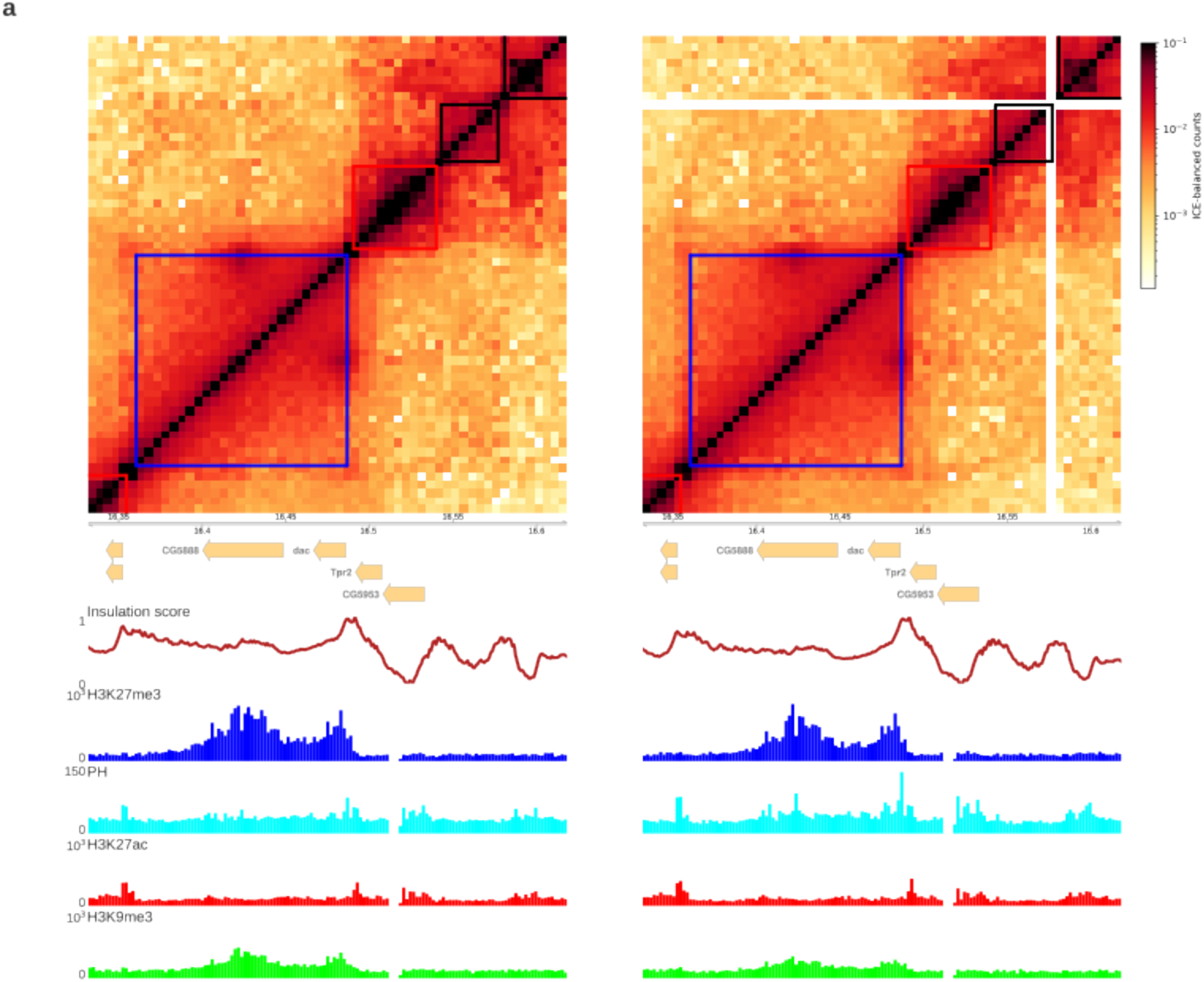
SUMO RNAi leads to genome-wide re-organization of the genome. **(a)** ICE normalised Hi-C maps for the genomic region around the *dac* locus in Control and SUMO RNAi samples. Colors towards black and red indicate high contact values, colors towards yellow and white indicate low contacts. C&R tracks for Pc, H3K27me3, H3K27ac, H2Aub118 marks in control and the SUMO RNAi sample shown in the bottom. Each square indicates a TAD called and the color indicates the enriched histone mark type: blue for H3K27me3- enriched, red for H3K27ac-enriched, and black for no enrichment of any used histone mark.

